# iPAR: A framework for modelling and inferring information about disease spread when the populations at risk are unknown

**DOI:** 10.1101/2024.11.08.622597

**Authors:** Stephen Catterall, Thibaud Porphyre, Glenn Marion

**Affiliations:** Biomathematics and Statistics Scotland, Edinburgh, United Kingdom; Laboratoire de Biométrie et Biologie Évolutive, UMR 5558, Universite Claude Bernard Lyon 1, CNRS, VetAgro Sup, Marcy l’Étoile, France

**Keywords:** epidemic modelling, host population, uncertainty

## Abstract

We introduce the inference for populations at risk (iPAR) framework which enables modelling and estimation of spatial disease dynamics in scenarios where the population at risk is unknown or poorly quantified. This framework addresses a gap in spatial infectious disease modelling approaches, with current methods typically requiring knowledge of the spatial distribution of the population at risk or making simplifying assumptions that are not supported in many real-world outbreaks where only case-report data are available. The principles for iPAR are demonstrated in the context of a susceptible-infected disease dynamics model coupled with Bayesian inference implemented via data-augmentation Markov chain Monte Carlo (MCMC). This implementation of iPAR is tested for a range of scenarios using simulated outbreak data. Results indicate that the method can effectively estimate key properties of disease spread from spatio-temporal case reports and make useful predictions of future spread. The method is then applied to a case study exploring the 2015-2019 Estonian outbreak of African Swine Fever (ASF) in wild boar. Estimates of epidemiological parameters reveal evidence for long distance transmission, as well as disease control via reduction of the wild boar population in Estonia.

## 1. Introduction

Outbreaks of infectious disease can be very costly and can have serious impacts on both natural systems and human economic activities (Knight-Jones and Rushton 2013; Kaye et al. 2021). For example, African swine fever (ASF), a disease that can infect both domestic pigs and wild boar, has spread since its introduction in Georgia in 2007 through many countries in Europe and Asia, leading to millions of culled pigs and consequent serious impacts on the global pig industry (Brown et al. 2021). More broadly, including for diseases of humans and plants, epidemic models can be usefully employed to assess risks arising from both current and future disease outbreaks (McBryde et al. 2020; Brooks-Pollock et al. 2015; Gilligan and Van Den Bosch 2008). Specifically, they can be fitted to reported outbreak data to infer important characteristics of disease spread (Swallow et al. 2022). These characteristics can then be used to inform potential disease control measures (Kretzschmar et al. 2022; Keeling et al. 2021).

The spatial distribution of the population at risk is very important because it strongly influences the contact structure of the population, which in turn is of key importance in the disease transmission process (Gudelj and White 2004). Further, spatially explicit epidemic models can offer insights and information critical to the control of real-world outbreaks e.g. identifying high risk areas and control zones (Boender et al. 2007; Porphyre et al. 2017). Typically, such models assume knowledge of the spatial distribution of the population at risk e.g. foot and mouth disease (FMD) transmission models (Keeling et al. 2001), Classical swine fever (CSF) transmission models (Gamado, Marion, and Porphyre 2017). However, a frequently encountered and challenging scenario is when we do not have complete knowledge of the population at risk of infection (Brooks-Pollock et al. 2015) and, in particular, we may not know the spatial distribution of this population. This is especially true for wildlife populations, which may not be well mapped, but can also frequently be the case for livestock. Records of livestock holdings may either be non-existent (especially for small ‘backyard’ holdings), out of date, incomplete or may not be publicly available for privacy reasons (e.g. farm locations in the USA (Tildesley and Ryan 2012)). In such cases there is a need to make use of case-only data. Here, we therefore focus on the scenario where cases are reported with possibly imprecise spatial and temporal coordinates.

A number of techniques have been proposed to model disease transmission when there is incomplete knowledge of the population at risk. One class of approaches involves modelling the disease case generating process, with the susceptible population not explicitly specified. Transmission trees are one example of this approach e.g. (Haydon et al. 2003). Another example is the use of techniques based on contact distribution models (Mollison 1977), for example (MacCalman et al. 2017; Lau et al. 2017). Such approaches benefit from the fact that, typically, more is known about disease cases than is known about general members of the susceptible population. On the other hand, this approach may be problematic in scenarios where the set of disease case reports is incomplete, or where depletion of susceptible individuals is an important factor in determining the progression of the epidemic. There is also a risk of bias in scenarios where individuals are highly mobile, such as susceptible wildlife populations. In such scenarios the location of a case (often where an animal is found dead) may be a poor proxy for the spatial history of the individual (Cross, Caillaud, and Heisey 2013). The above referenced approaches do not allow for spatial variation in susceptible population density, with the exception of (Lau et al. 2017) in which the susceptible population density is both allowed for and known.

A second class of approaches, which has been applied to livestock epidemic data, involves making assumptions/predictions about unknown farm locations to permit spatially explicit transmission modelling. For example, assuming that farms are located uniformly at random across the landscape can sometimes be sufficient to identify optimal control measures (Tildesley et al. 2010), though this can only be applied to certain controls (ring culling) and requires extensive outbreak data. However, using land use data and other landscape covariates to predict farm locations can improve the accuracy of livestock epidemic models (Sellman et al. 2020; Tildesley and Ryan 2012). Such approaches seem useful in scenarios where we know some aspects of the susceptible population, e.g. for livestock the total number of farms in the region of interest, but it is unclear how they might be applied to less well mapped populations such as wildlife, ‘backyard’ holdings or in jurisdictions where data is more limited.

We consider how to model disease spread when the susceptible population density is unknown, but we have spatial covariates that may explain population density and other aspects of the population of relevance to the transmission process. Intuitively, case reports with approximate spatial locations are informative of the distribution of the population at risk, and coupled with observation times also provide insight into rates of spread in space and time. To motivate our proposed approach, consider again how knowledge of the population at risk is used to construct spatially explicit epidemic models, e.g. those used to model the spread of FMD (Chis Ster, Singh, and Ferguson 2009). The rate of transmission between an infectious location and a susceptible location is typically assumed to be an overall rate multiplied by (a) a kernel function depending on the distance between the locations, (b) the infectivity of the infectious location and (c) the susceptibility of the susceptible location. Thus, from a modelling perspective, in the most general setting, knowledge of the population at risk is captured in two ‘surfaces’ covering the landscape: an infectivity surface and a susceptibility surface. In reality, although these surfaces will be influenced by the spatial distribution of the population at risk, they are also influenced by many other factors e.g. variations in biosecurity, variations in host susceptibility. In this paper, assuming limited or no knowledge of the population at risk, we develop an approach to estimate these two surfaces along with key characteristics of spatial disease transmission, given reported disease cases, but avoiding making any explicit assumptions about the spatial distribution of the population at risk. We refer to this as the inferring - information about - populations at risk (iPAR) framework.

To constrain the set of possible surfaces, we assume that each surface is a function of multiple landscape covariates, with parameters estimated from the disease case data. We also aggregate spatially to a set of patches covering the landscape, with the patches being the infectious units of the model. This aids computational tractability but also enables use of uncertain spatial locations. Patch-based models have been used successfully to model the spread of infectious disease, the spread of invasive non-native plant species and other spatio-temporal ecological processes (Lau et al. 2014; van den Driessche 2008; Catterall et al. 2012). Although spatial aggregation can result in a loss of spatial resolution, the degree of aggregation (patch size) can be controlled to minimise any potential loss of information. The iPAR framework is also highly flexible and can make use of wide array of spatial covariate data. If suitable covariates are available, we can expect that at least some of the variation in patch susceptibility/infectivity can be explained by the covariates. Land use and climate data could, thus, be useful as covariates in most situations. Other covariates could be useful in specific scenarios e.g. human population density, hunting bag data and previously published estimates of the susceptible population density.

In this paper we describe an initial implementation of the iPAR framework outlined above. This is described in detail (Section 2.1) along with the MCMC implementation of Bayesian inference (Section 2.2) used for parameterisation from disease case reports for the region of interest. This inference framework is extensively tested using simulated case data (Section 3). We illustrate the potential of the iPAR framework through application to case reports arising from the ASF outbreaks in Estonia that occurred between 2014 and 2019 (Section 4). This analysis reveals evidence for both long distance transmission and the effectiveness of disease control programmes in Estonia. The case study demonstrates the value of the iPAR approach, and we discuss potential application to other systems and possible future development of the iPAR framework (Section 5).

## 2. The iPAR framework

Here we define an underlying spatial disease transmission model and associated inference tools that together provide a flexible approach to deal with the problem of missing information on populations at risk in spatial epidemiology. This defines an initial implementation but note that many extensions are possible within the iPAR framework.

### 2.1 The generic model

We model transmission of disease in continuous time across a region *R*. We partition the region *R* into a collection of *I* patches, with each patch indexed by an integer *i* (1 ≤ *i* ≤ *I*) and associated with a given sub-region of *R*. Within the region *R*, the spatial distribution of the population at risk is uncertain. However, we assume that we have covariates available that can be used to model the variation in patch susceptibility/infectivity across the region. The chosen size of the patches depends on a number of factors, including the spatial resolution of the covariates and the disease case data, as well as the scale of apparent disease spread in the region. These patches are the units of infection within the model framework. For each patch *i*, there is a set of associated covariate values 𝒞_*i*_. In principle, the covariates may vary both temporally and spatially. However, for notational simplicity, we do not show explicit time dependence in the presentation that follows, and in all the applications of the methodology that we consider in this manuscript the covariates are constant in time.

For simplicity we consider modelling transmission according to a spatial SI model. That is, each patch is either susceptible to infection or has become infected/infectious. However, we note in general that more complex dynamics and a greater set of possible states could in principle be accounted for if suitable data were available. The SI assumption is most appropriate for the early stages of a disease outbreak, but can continue to be reasonable if the disease is persistent at the patch level on the timescales of the outbreak and this criterion should be considered when selecting an appropriate patch size for the analysis – for example, see the case study in Section 4 below. The transmission of disease is modelled over some time interval [0, *T*] via a continuous time discrete state space Markov process. The initial conditions comprise a list of the patches that are infected at time *t* = 0, with the remainder assumed susceptible.

Given the initial conditions, the transmission model is fully specified once we define the force of infection on a susceptible patch at any point in time. The force of infection

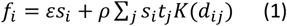

on a susceptible patch *i* is comprised of two terms, the first term representing primary infection due to transmission from outside the region under study, the second term representing transmission from infected patches within the region *R*. Each patch *i* has susceptibility *s*_*i*_ and, when infected, infectivity *t*_*i*_ which can be functions of spatially varying covariates – meaning that they vary between patches.

The parameter *ε* controls the overall rate of background infection, while *ρ* controls the overall rate of secondary transmission. The summation is over all infectious patches *j, K* is a so-called transmission kernel and *d*_*ij*_ is the distance between the centroids of patches *i* and *j*. Typically, we use a truncated power law kernel *K*(*d*) = *d*^−2*λ*^*I*(*d* < *d*_max_)/*m*(*λ*) where *m*(*λ*) is a normalisation constant defined in the ESM (Appendix 2), *d*_max_ is a fixed maximum transmission distance, and *λ* is a parameter estimated from the outbreak data. If the patches are all squares of the same size then distance, as an input to the kernel function, is measured in units such that one unit is equal to the width of a patch, meaning that e.g. *d*_*ij*_ = 1 precisely when patches *i* and *j* are nearest neighbours. The above expressions together represent what we call the *constant-in-time* model, although as noted above this can accommodate temporal variation in the covariates.

We also consider a variant of this model, which we term the *varying-in-time* model, in which the total force of infection is multiplied by a function of time *h*(*t*). This function is intended to model the effects of disease control. For identifiability we assume that *h*(0) = 1, with *h* then typically decreasing following the application of control measures such as removal of hosts. However, we assume only that *h*(*t*) > 0 for all *t*, allowing *h*(*t*) to both increase and decrease over time, so that such changes are estimated from the data.

We now define the susceptibility *S*_*i*_ and infectivity *I*_*i*_ of a patch *i* in terms of the covariates for that patch. There are many ways of defining the relationship between susceptibility/infectivity and the covariates. Here we suggest a general approach. Typically, covariates can carry either absolute or relative information. Absolute covariates might include, for example, average temperature and rainfall for the patch. Relative covariates might include, for example, the proportion of the patch occupied by a specific land use such as forest. Suppose that the set of covariates *C*_*i*_ for patch *i* comprises absolute covariates *c*_*i,k*_, 1 ≤ *k* ≤ *K* and relative covariates *h*_*i,l*_, 1 ≤ *l* ≤ *L*, with *h*_*il*_ being the proportion of patch *i* occupied by land use category *l* (derived from a partition of land into *L* distinct land uses). Then, a plausible model for the susceptibility of a patch is 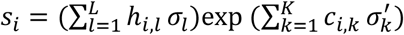 where the Greek letters are parameters to be estimated and each *σ*_*l*_ ≥ 0. We also require that 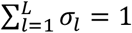 for identifiability, bearing in mind that *s*_*i*_ will be multiplied by the overall rate *ρ* in the expression for the secondary force of infection.

Each land use category contributes additively to the susceptibility, with the contributions weighted by the covariates *h*_*i,l*_. Each absolute covariate contributes multiplicatively, with negative relationships possible for these covariates when 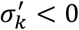. However, it is important to note that negative relationships are also possible for the land use covariates, even though the *σ*_*l*_ are non-negative, due to the constraint that 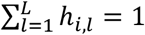. For example, the covariate *h*_*i,l*_ with the smallest associated *σ*_*l*_ will be negatively related to susceptibility. This general form of susceptibility function is analogous to the suitability function defined in (A. Cook et al. 2007) in the context of biological invasions. A similar expression is assumed for patch infectivity 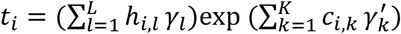. Again, the Greek letters 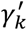 for 1 ≤ *k* ≤ *K* and *γ*_*l*_ for 1 ≤ *l* ≤ *L* are parameters to be estimated, with each *γ*_*l*_ ≥ 0 and 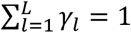. This completes the specification of the generic model.

### 2.2 Bayesian inference and prediction based on disease case reports

#### 2.2.1 Parameter and latent variable estimation

Having defined the model, we now present methodology that allows fitting of the model to data including estimation of unobserved (latent) infection times. For a disease outbreak, case reports describe observed cases of the disease in individual animals or on individual farms. We assume that each case has a location and a detection time associated with it. Locations are taken to be specified at patch resolution or better. Detection times of disease cases are taken to be interval-censored, which allows for the common scenario whereby new case reports are provided at regular intervals (e.g. daily, or monthly), but can also accommodate irregular observation times. We denote these reporting times by 0 = *t*_0_ < *t*_1_ < ⋯ < *t*_*N*_ = *T* where [0, *T*] is the time interval over which case report data are available. We assume that interval-censored detection times should constrain the actual infection times of the individuals/farms within the patch i.e. that a case occurring after time *t*_*i*_ but before time *t*_*i*+1_ is reported at time *t*_*i*+1_. In practice, we expect performance of this modelling framework to degrade if a significant number of infections occurring in an interval (*t*_*i*_, *t*_*i*+1_) are unobserved at *t*_*i*+1_. To some extent this can be countered by extending these reporting time windows used in the analysis resulting in greater estimated uncertainty in inferred infection times (see below). When aggregating to the patch level, we define the infection time of a patch to be the earliest recorded infection time of the set of individuals/farms within it.

We use Bayesian techniques to infer the model parameters from outbreak data. The true infection time of each patch is not known with certainty so, as is common when fitting epidemic models, we use data augmented MCMC to facilitate the inference (Swallow et al. 2022). This general approach to inference is well established in the literature and has been applied in, for example, (O’Neill and Roberts 1999), (Jewell et al. 2009) and (Stockdale, Kypraios, and O’Neill 2017). Let *θ* denote the set of model parameters and let 𝒰 = {*U*_*i*_; 1 ≤ *i* ≤ *I*} denote the set of unobserved patch infection times. For each patch *i*, let *T*_*i*_*ϵ*{0,1,2, …, *N* + 1} denote the ‘time category’ into which the patch falls in terms of reporting. So, *T*_*i*_ = 0 if *i* was already reported as infected at time *t*_0_ = 0 and *T*_*i*_ = *j* (1 ≤ *j* ≤ *N*) if *i* was first reported as infected at reporting time *t*_*j*_. Finally, *T*_*i*_ = *N* + 1 if *i* was not reported as infected at any point in the time interval [0, *T*], which we assume implies that *i* remains susceptible at time *T*. Finally, let 𝒯 denote the set of observed *T*_*i*_. These observed time categories 𝒯 are the data we use to infer the model parameters *θ* and the actual and unobserved or latent patch infection times 𝒰. Bayes’ theorem states that

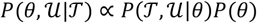

Here *P*(*θ*) represents the joint prior distribution of the model parameters. The prior distribution is typically chosen to be non-informative (see ESM Appendix 3 for details). The factor *P*(𝒯, 𝒰|*θ*) above is usually termed the complete data likelihood for the model parameters conditioned on the observed data and a given set of latent infection times. The transmission process is a continuous time discrete state space Markov process, so this likelihood can be computed using standard techniques as described in, for example, (Gibson and Renshaw 1998; O’Neill and Roberts 1999). We obtain, for the varying-in-time model,

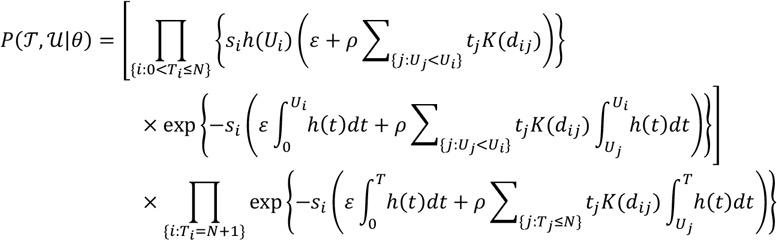

The above expression is conditioned on the history of transmission prior to time *t* = 0 and written under the assumption that the time category observations 𝒯 are consistent with the given infection times 𝒰. The second product represents the probability that patches never observed to be infected escape infection up to time times *T*. On the other hand, the first product is over patches observed as infected in the time interval [0, *T*] and represents the probability of their infection at the latent infection times 𝒰. Proposed changes to 𝒰 - made in the sampling algorithm described in the ESM (Appendix 1) - must therefore be consistent with 𝒯. For computational simplicity we assume that *h* is piecewise constant on pre-defined subintervals of [0, *T*] with *h*(0) = 1 for identifiability, then *h*(*t*) = *h*_*i*−1_ on the *i*’th subinterval (2 ≤ *i* ≤ *H*). The values *h*_*i*−1_ are regarded as parameters to be estimated from the data and they are appended to the parameter set *θ*.The corresponding likelihood for the constant-in-time model is obtained by setting the function *h* equal to 1 for all times *t*.

Having obtained an expression for the complete data likelihood, we are now in a position to draw samples from the joint posterior distribution *P*(*θ*, 𝒰|𝒯). This is achieved by Markov Chain Monte Carlo (MCMC) methods. Full details of the sampling algorithm may be found in the ESM (Appendix 1).

#### 2.2.2 Model selection

Model selection is a challenging problem, and is not the main focus of this manuscript. One widely used metric for assessing goodness of fit is the Deviance Information Criterion (DIC) (Spiegelhalter et al. 2002). The DIC is not uniquely defined for latent variable models (Celeux et al. 2006). In (Gamado, Marion, and Porphyre 2017) they assessed a number of DIC variants in the context of spatio-temporal epidemic modelling and found that their metric *DIC*_1_ (which corresponds to *DIC*_6_ in (Celeux et al. 2006)) performed well. Therefore we use this metric to assess models in the current manuscript, in particular to compare the constant-in-time and varying-in-time models when both fitted to the same set of outbreak data. See ESM (Appendix 4) for details of the definition and computation of this metric.

#### 2.2.3 Uncertainty in model predictions

Having fitted the model to outbreak data, we can then use the fitted model to predict the timing of future transmission events beyond the period of the observed data by sampling from the appropriate posterior predictive distribution. This is achieved by forward simulation of the transmission model over a time interval [*T, T* + *S*], with model parameters sampled from the joint posterior distribution and initial conditions typically set to the infection status of the patches at the final time point of the data used to fit the model (*t* = *T*). We typically conduct large numbers of such simulations, at least 10000 say, so as to capture the full distribution of possible outcomes.

The model predictions obtained from sampling the posterior predictive distribution can be summarised in a number of ways. We can plot the number of infected patches as a function of time. We can also plot risk maps that depict, for each patch *i*, the probability that it is infected by some future time *t* = *T* + *S*, which we calculate as the proportion of simulations in which *i* becomes infected by time *t* = *T* + *S*.

The use of the constant-in-time model to predict future events in this way is straightforward. On the other hand, fitting the varying-in-time model leads to an estimate of the function *h*(*t*) for *t* < *T*, but we cannot use this fitted model to make predictions without further assumptions on the behaviour of *h*(*t*) for *t* > *T*. For this reason, we only use the constant-in-time model to make predictions, although one could envisage future projections conditioned on specified scenarios for *h*(*t*) at *t* > *T*.

The predictions can also be compared to what actually happened, if such data are available. This is possible for simulated data where we generate a realisation of the model that represents the ‘truth’. Alternatively, with real data from a disease outbreak we can hold out some portion of the data from later stages of the outbreak, fitting the model only to data from the earlier stages of the outbreak. In either case, we can compare the predictions to such ‘truth’ in two ways. First, we may directly compare the *total number* of infected patches as it increases over time. Second, we can assess the ability of the predictions to identify the *correct* infected patches, rather than simply the correct overall number of infected patches, by computing *discrimination* metrics that compare the risk map to the true map of infection. For a given probability threshold *p*, we calculate the true positive rate (TPR, the proportion of patches infected during [*T, T* + *S*] with infection probability exceeding *p*) and the false positive rate (FPR, the proportion of patches uninfected at time *t* = *T* + *S* with infection probability exceeding *p*). Plotting the true and false positive rates for multiple thresholds *p* gives the Receiver Operating Characteristic (ROC) curve (Fielding and Bell 1997), from which standard metrics such as the AUC (Area under the ROC curve) and the TPR at 5% FPR can be derived.

#### 2.2.4 Testing the iPAR modelling approach

To assess the iPAR framework, it is important to first test it using simulated outbreak data. The use of simulated data has two key benefits. First, we can explore a wider range of scenarios than is possible using real outbreak data, which can be very helpful when exploring the strengths and weaknesses of methods, as well as the limits of their applicability. Second, with simulated data we always know the ‘true’ data generating process, which allows us to assess, for example, how close parameter estimates obtained from the simulated data are to their ‘true’ values. Such comparisons between true and estimated parameter values are impossible with real outbreak data. Nonetheless, it is also critical to test methods via application to real world data and problems. Model assumptions can be validated externally by comparing fitted model outputs with other sources of data on the outbreak and the susceptible population that were not used to fit the model. Such approaches depend very much on the details of the scenario under investigation. We apply the iPAR framework to disease case report data and describe some specific validation approaches in the case study in Section 4.

Simulation-based methods are used to systematically test the constant-in-time iPAR model in Section 3.2. We employ a form of sensitivity analysis in which we start from a baseline model parameterisation based on the parameter estimates obtained from the ASF outbreak data in the case study in Section 4. We then vary each model parameter separately (keeping all others fixed), simulating a single set of outbreak data for each parameter combination and then inferring model parameters from the simulated outbreak data. In this way, we can build up plots of true versus inferred parameter values (95% credible intervals) for each model parameter. This kind of sensitivity analysis has been conducted in e.g. (Pooley et al. 2020). If the model parameters are sampled from the prior then the *average* coverage is equal to the nominal value (S. R. Cook, Gelman, and Rubin 2006) but, for a specific choice of parameters, Bayesian credible intervals don’t necessarily have their nominal coverage. This means that the true value of a parameter may not fall within a 95% Bayesian credible interval for 95% of the time. Nevertheless, we can visually assess the output from the sensitivity analysis in order to check (a) whether the inference is recovering the ‘true’ parameter values in the way we would expect, and (b) which parameters are more difficult to recover from the simulated data. For parameters that are difficult to recover, the inferred parameter may only depend weakly on the true parameter, and may instead be strongly influenced by the prior distribution assigned to the parameter.

In the setting of the constant-in-time model with no absolute covariates, which is the setting we use in Section 3.2 below, the full set of model parameters is as follows: the land use susceptibilities *σ*_1_, …, *σ*_*L*_, the land use infectivities *γ*_1_, …, *γ*_*L*_, and three remaining parameters *λ* (kernel parameter), *ρ* (overall transmission rate) and *ε* (background transmission rate). In order to perform the sensitivity analysis, each parameter is taken in turn and varied while keeping other parameters fixed. One issue is that the susceptibilities are not independent, being constrained to sum to 1, and similarly for the infectivities. To circumvent this problem, we reparametrize the model using the isometric logratio transform *ξ* = ilr(σ)∈ℝ^*L*−1^, *η* = ilr(γ)∈ℝ^*L*−1^, which maps the simplex-valued parameters to real multivariate parameters (see ESM Appendix 5 for details). Thus, the components of *ξ* and *η* are unconstrained and can be varied independently as part of the sensitivity analysis.

## 3. Assessing the iPAR framework performance

We now assess the iPAR framework by applying it to simulated outbreak data. In Section 3.1, we analyse data from a single simulated outbreak – simulated from the constant-in-time model with spatially varying susceptibility/infectivity – in order to illustrate the typical workflow and outputs obtained from the model fitting process. In Section 3.2, we conduct a systematic simulation study in which we generate outbreak data for a wider range of scenarios, though still for the constant-in-time model with spatially varying susceptibility/infectivity, and assess the ability of the inference to recover the ‘true’ parameters. In Section 3.3 we explore the ability of the inference to recover temporal trends in transmission, using simulated outbreak data from the varying-in-time model with spatially varying susceptibility/infectivity. Finally, in Section 3.4, we assess the benefits of modelling spatial variation in susceptibility and infectivity, one of the key aspects of the iPAR framework, both in terms of estimating key parameters and in terms of predicting held-out future simulated outbreak data. To do this, we generate simulated outbreak data from the constant-in-time model with spatially varying susceptibility/infectivity, but we fit either a model of similar form or a model which assumes spatially homogeneous susceptibility/infectivity.

### 3.1 Estimation and prediction for an illustrative simulated outbreak

Outbreak data are simulated using the constant-in-time iPAR model on the Estonian landscape, which is divided into 575 10km square patches, with model parameters similar to those estimated from the outbreak data for ASF in wild boar in Estonia (see the case study in Section 4). As in the case study, land use data for Estonia are used as the basis for computing covariates used to model spatial variation in both susceptibility and infectivity. Four patches are initially seeded with infection, and the duration of the simulation is chosen such that a substantial proportion of patches become infected by the end of the simulation.

A single simulated outbreak can be summarised by plotting the number of infected patches over time (Figure 1A, black line). The constant-in-time iPAR model was fitted to the early-stage epidemic data up to month 12 from this single simulated outbreak, with simulated patch infection times interval censored into one-month long intervals. The resultant posterior distribution provides estimates for all model parameters. For example, we can visualise the posterior marginal distribution of the background transmission rate parameter *ε* (Figure 1B, with true value represented by a vertical line) by aggregating MCMC samples. We can also generate the 95% credible intervals for parameters and functions of the model parameters such as the transmission kernel (Figure 1C, true kernel in black, 95% credible interval in blue shading). Credible intervals for the components of the susceptibility parameter *σ* (Figure 1D, 95% credible intervals represented by blue lines) are often quite wide, but in general these intervals align well with the ‘true’ values.

**Figure 1.**
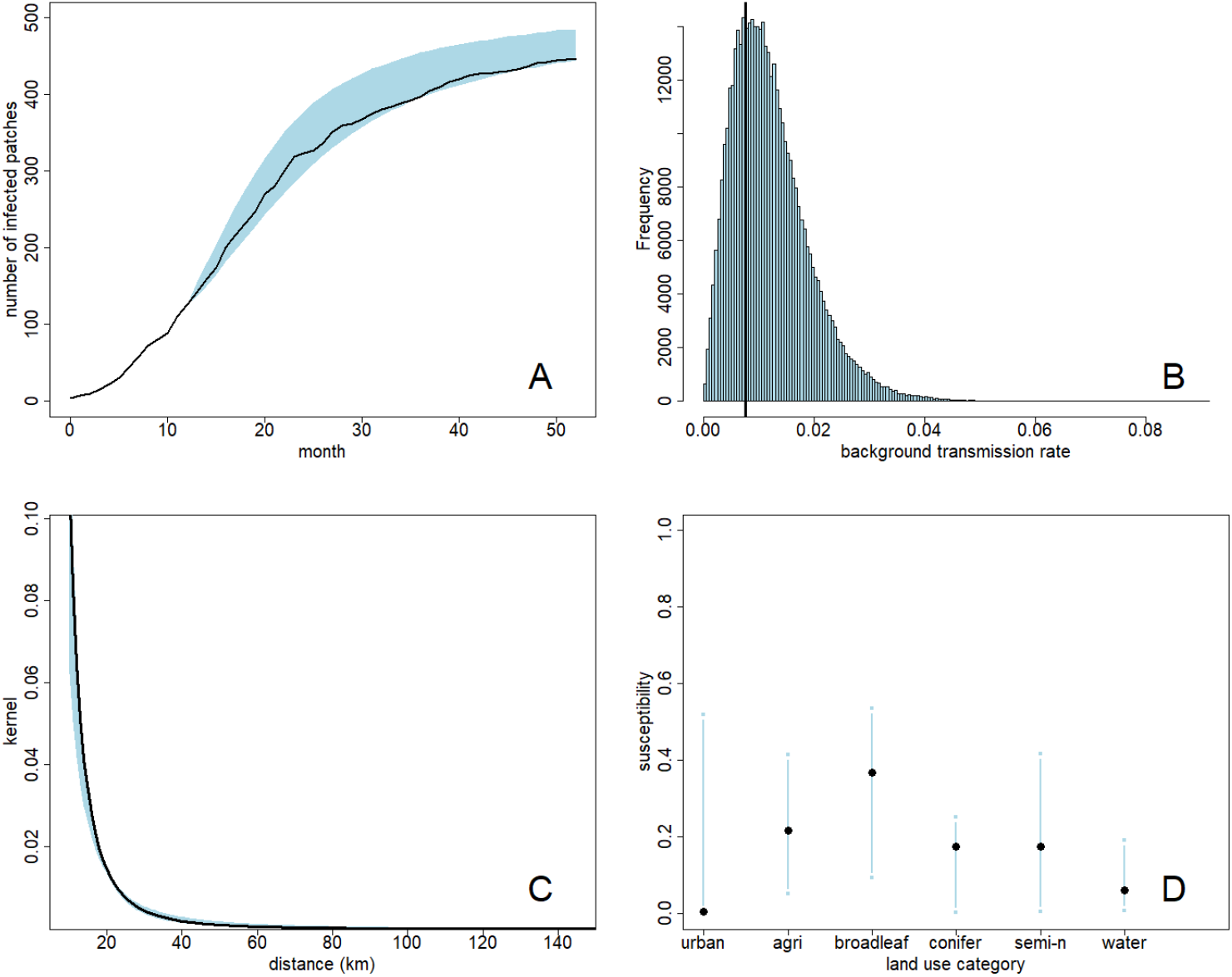
Estimated parameters and predictions from fitting the iPAR model to simulated outbreak data. A: number of infected patches over time in the simulated outbreak data (black line). The model was fitted to the outbreak data up to month 12. Predicted number of infected patches (95% credible interval) beyond month 12 is shown in blue. B: marginal posterior distribution for the background transmission rate, with ‘true’ value indicated by the black vertical line. C: ‘true’ kernel function (black line) and estimated kernel function 95% credible interval (blue shading). D: estimated susceptibility parameter *σ* for each land use (95% credible interval, blue) and ‘true’ value (black circle).

The model, fitted to data up to month 12, was then used to predict progression of the epidemic beyond the first 12 months. This was achieved by simulating 10000 times with parameters sampled from the posterior distribution and initial conditions specified by the state of the epidemic at month 12. These predictions may be summarised by 95% credible intervals for the number of infected patches over time (Figure 1A, blue shading). The intervals contain the ‘true’ epidemic curve in this case, suggesting satisfactory inference of the model parameters.

### 3.2 Estimation of key epidemiological parameters

We systematically tested the constant-in-time model over a wide range of parameter combinations using the sensitivity analysis approach described in Section 2.2.4. The simulation model set-up was identical to that in Section 3.1 except for the use of different sets of model parameters. The data from most of the simulated outbreak (up to month 52) was used to fit the model. Plots of true versus inferred parameters are available in Figure 2 for *λ, ρ, ε* and in the ESM (Appendix 6) for the remaining parameters. Key model parameters such as the kernel parameter *λ* and the background transmission rate *ε* are well recovered from the simulated outbreak data, though the overall transmission rate *ρ* tends to be somewhat overestimated. Reasons for the latter phenomenon will be discussed later. The susceptibility parameters tend to be well recovered, while the infectivity parameters seem more difficult to recover, which is consistent with the results in (Pooley et al. 2020).

**Figure 2.**
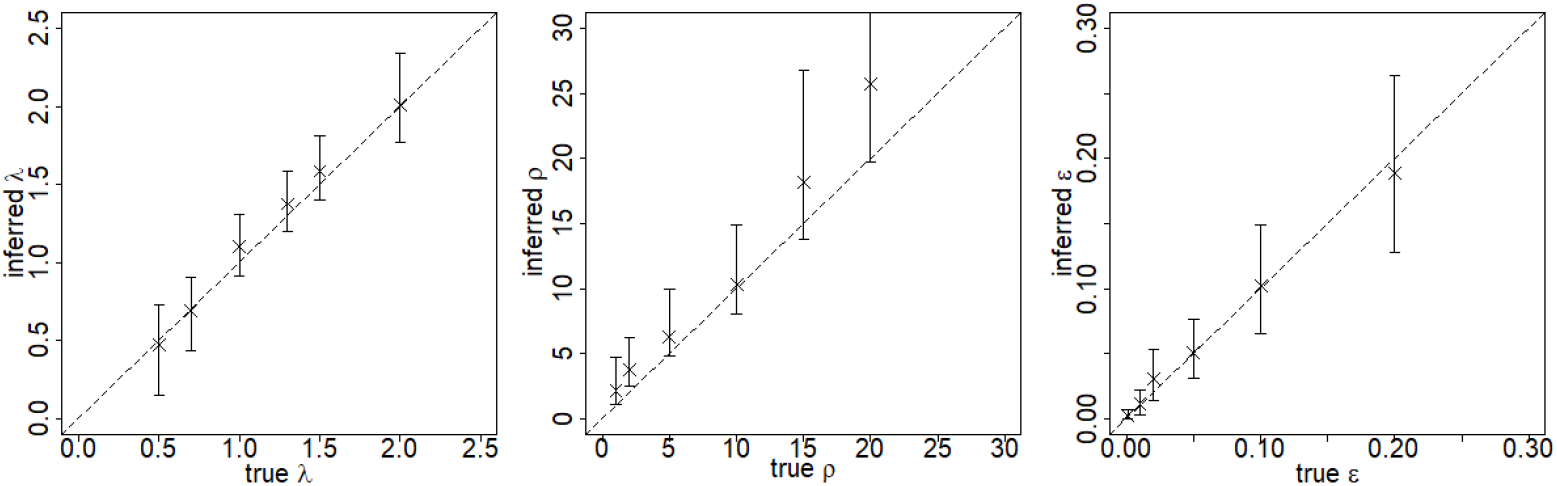
Assessment of the reliability of parameter inference for the constant-in-time iPAR model. Each panel corresponds to a parameter of the model. The ‘true’ value of the parameter is plotted against the posterior median (cross) and posterior 95% credible interval (vertical line). For comparison purposes we also superimpose the diagonal line representing perfect agreement between inferred and true values. This figure includes panels for the kernel parameter *λ*, the overall transmission rate *ρ* and the background transmission rate *ε*. Analogous figures for the other parameters may be found in the ESM (Appendix 6).

### 3.3 Estimation of temporal trends in transmission

Next, we focus on the varying-in-time iPAR model with parameters similar to those obtained from fitting this variant of the model to the Estonian ASF outbreak data (see case study in Section 4) as a baseline parameterisation. We then vary the form of the temporal trend function *h* according to three pre-chosen scenarios: (1) *h* decreasing linearly (perhaps representing increasing levels of disease control), (2) *h* constant (no disease control), and (3) *h* initially constant then suddenly drops to low levels (corresponding to rapid strong disease control after some delay). The changepoints of the function are fixed and are kept equal to those used in the case study. The function *h* from the first scenario is closest to what was actually estimated from the data in the case study (Table 4). Note that, in the case study, *h* was seen to initially increase before decreasing, which is why we have also allowed for an initial increase in *h* in the first and third scenarios. We assess the extent to which these temporal trends can be recovered by inference from data simulated under each of the above three scenarios. We also fit the constant-in-time model to the same sets of simulated outbreak data, which allows us to compare model fits (constant-in-time versus varying-in-time) using *DIC*_1_ as discussed in Section 2.2.2.

Figure 3 shows the simulated ‘true’ incidence curve for each of the three temporal trends. Overlaid on this ‘true’ incidence curve is a region representing 95% credible intervals for incidence, obtained from simulations of the fitted varying-in-time model with initial conditions the same as those of the simulated outbreak data used for fitting the model. For all three temporal trends, the agreement between the credible intervals and the ‘true’ incidence curve shows that the algorithm is able to reliably recover parameters giving temporal trends in incidence that agree with the ‘true’ incidence curve. Figure 3 also shows the 95% credible interval for *ρh*(*t*) in the same set of three simulations. Despite the striking fact that the form of this function is not at all obvious if presented with a single outbreak incidence curve, the trend function *ρh*(*t*) is inferred correctly from single outbreaks, though with some overestimation. We also computed *DIC*_1_ values for each model fit (see ESM Appendix 4 for actual values obtained). For scenarios (1) and (3), in which *h* ≠ 1, the *DIC*_1_ indicated a strong preference for the varying-in-time model. For scenario (2), in which *h* = 1, there was a mild preference for the constant-in-time model. These results were all as we would expect.

**Figure 3.**
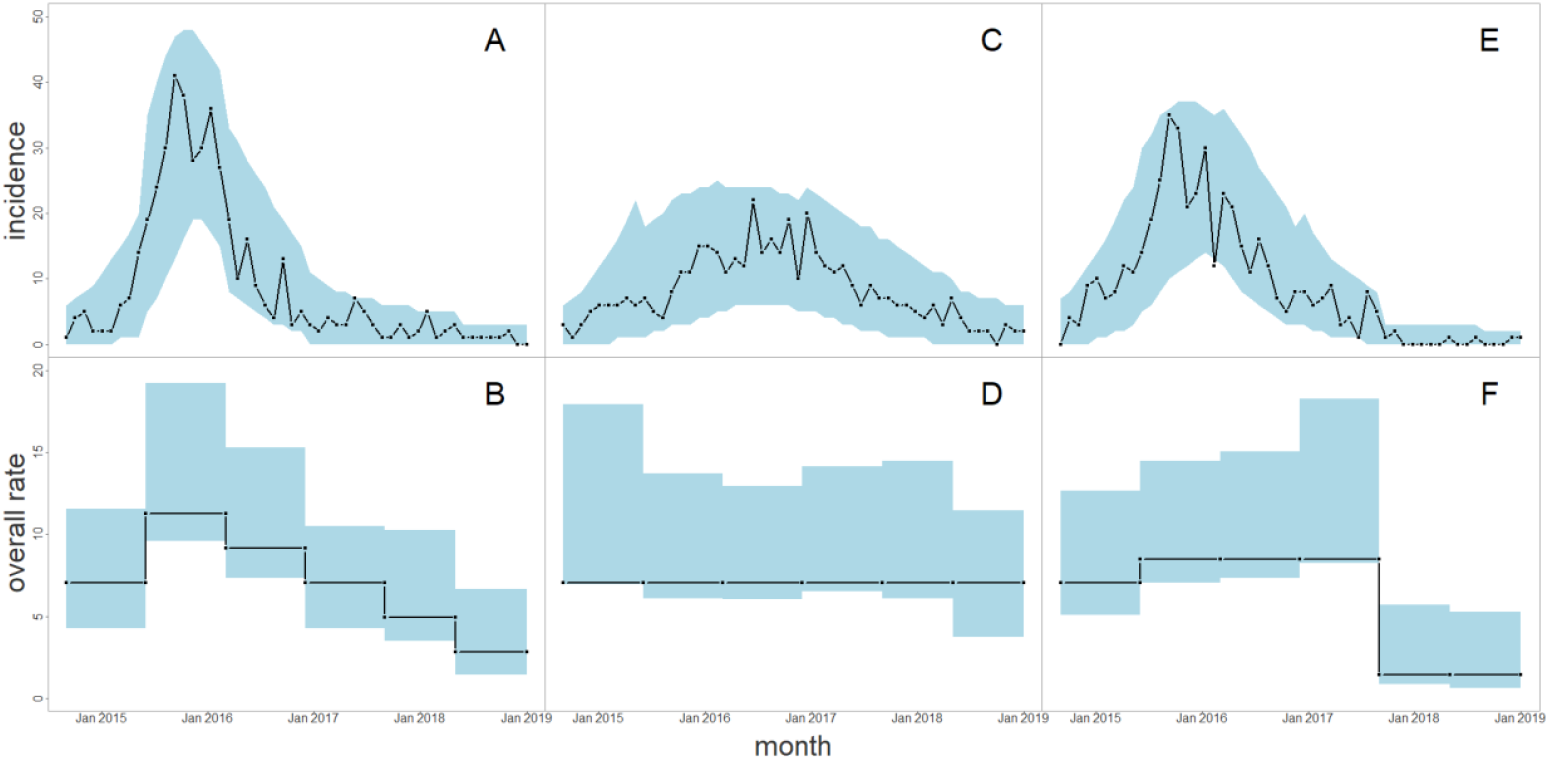
Incidence curves and corresponding estimates of the overall rate *ρh*(*t*) from the time varying simulation study. A: the black line shows the incidence curve from a single simulation of the “linear decrease” scenario. The blue area shows the 95% credible intervals for incidence, defined as newly infected patches each month, obtained from the posterior predictive distribution of the fitted varying-in-time model, given the data from that single simulation. B: true and estimated *ρh*(*t*) for the same scenario. Black line represents true value, blue area shows the 95% credible interval. C,D: same plots for the “constant over time” scenario. E,F: same plots for the “sudden drop off” scenario (see text for details).

### 3.4 Benefits of modelling spatial variation in susceptibility and infectivity

We assess the importance of modelling spatial variation in susceptibility and infectivity in the context of the constant-in-time iPAR model with land use covariates but no absolute covariates. The spatial variation in the patch susceptibilities and infectivities of the model arises as a result of spatial variation in the covariates, combined with differences in the susceptibility and infectivity parameters between land use categories. Our approach here is to simulate outbreak data incorporating such spatial variation and then fit two model variants to the simulated outbreak data. First, we fit a heterogenous model which accounts for this spatial variation and has the same form as the model used for simulation. Second, we fit a simpler homogeneous model in which there are no covariates and patch susceptibility/infectivity is spatially constant. Comparing the two model fits and their associated predictions enables us to assess the importance of taking into account variation in susceptibility and infectivity.

We consider model fitting given two levels of data availability. First, we fit models to data from the early stages of an outbreak. Second, we fit models to data from an entire outbreak. At both levels, we consider four different parameter combinations for the simulated outbreak data. This is less than the number of parameter combinations considered in Section 3.2, however the chosen parameter combinations cover a range of common transmission patterns. We use either a wide or a moderately wide transmission kernel. Narrow kernels, such as those approximating nearest neighbour kernels, are avoided because we intuitively expect there to be little difference between the predictive performance of the heterogenous and homogeneous models in this case – see the case study in Section 4 and associated discussion for more on this topic. In the simulations, we assume that there is a single land use with high susceptibility (all other land uses are assigned equal low susceptibilities), and the same applies for infectivity. Broadleaf woodland is taken to have high susceptibility in all scenarios, while the high infectivity land use alternates between broadleaf woodland and arable land. Precise details of the scenarios are provided in the ESM (Appendix 8).

We repeat the model fitting and prediction for 20 replicate simulations per parameter combination, in order to give a fuller picture of parameter recovery and predictive performance in each case. Mean (over all 20 replicates) AUC and mean TPR at 5% FPR measure discriminatory power. We compute the posterior predictive distribution of *N*_*est*_/*N*_*truth*_, the predicted number of infected patches divided by the true number of infected patches, and its reliability which is the % of replicates in which the 95% CrI for *N*_*est*_/*N*_*truth*_ includes *N*_*est*_/*N*_*truth*_ = 1. We also compute the mean bias (posterior median of *N*_*est*_/*N*_*truth*_minus 1, averaged over all 20 replicates). Corresponding quantities are computed for the kernel parameter *λ*. For the susceptibility and infectivity parameters we adopt a different approach. We focus on estimation of the large susceptibility parameter and the large infectivity parameter. These parameters are likely more difficult to estimate than the other susceptibility/infectivity parameters because they are far from the main mass of the prior distribution on the simplex. So, we measure the difference between the posterior and prior medians (averaged across all 20 replicates), which captures the extent that the posterior has moved away from the prior. If the difference is very small then there is little information in the data regarding that parameter. For the given simulation set up, a posterior median that is 0.73 higher than the prior median corresponds to perfect estimation of the large susceptibility/infectivity parameter.

Using only early stage data, predictive performance assessed using discrimination metrics was consistently better for the heterogeneous model, if not by a very large margin (Table 1). Ability to predict *N*_*est*_/*N*_*truth*_ was good for both the heterogenous and homogeneous models. In terms of parameter recovery using only early stages data, we focus on the transmission kernel parameter, the susceptibility parameter associated with the high susceptibility land use class and the infectivity parameter associated with the high infectivity land use class. The kernel parameter was typically well estimated (Table 2), though the homogeneous model tended to somewhat overestimate this parameter, a phenomenon that was also observed by (A. Cook et al. 2007). However, susceptibility and infectivity parameters were imperfectly recovered (Table 2). Comparison of the posterior and prior medians suggests that the early stage data used for fitting provides some information on susceptibility but is less informative on infectivity.

**Table 1.** Early stage data predictive performance. Models are fitted to data on the early stages of simulated outbreaks on the Estonian landscape. Twenty independent replicates were carried out for each of the four parameter combinations. For each parameter combination (choice of transmission kernel and choice of high susceptibility and high infectivity land uses), we compare predictions from fitted homogeneous and heterogeneous models at four months beyond the final month of the simulated outbreak data used to fit the model.

| Scenario | | | | Discrimination metrics | | $N_{est}/N_{truth}$ | |
| --- | --- | --- | --- | --- | --- | --- | --- |
| Kernel | Susc | Infe | Model | mean AUC | mean TPR @ 5% FPR | reliability (%) | mean bias |
| wide | broadleaf | broadleaf | homog | 0.67 | 0.13 | 85 | -0.02 |
|  |  |  | heter | 0.71 | 0.18 | 90 | -0.03 |
| wide | broadleaf | arable | homog | 0.70 | 0.15 | 95 | 0.00 |
|  |  |  | heter | 0.74 | 0.20 | 95 | -0.01 |
| moderate | broadleaf | broadleaf | homog | 0.74 | 0.18 | 95 | 0.02 |
|  |  |  | heter | 0.78 | 0.25 | 100 | -0.00 |
| moderate | broadleaf | arable | homog | 0.74 | 0.19 | 85 | 0.04 |
|  |  |  | heter | 0.78 | 0.26 | 90 | 0.01 |

**Table 2.** Early stage data parameter recovery. Models are fitted to data on the early stages of simulated outbreaks on the Estonian landscape. Twenty independent replicates were carried out for each of the four parameter combinations. Mean post-prior is the mean, over replicates of the posterior median minus the prior median for the parameter.

| Scenario |  |  |  | Kernel parameter |  | Susceptibility | Infectivity |
| --- | --- | --- | --- | --- | --- | --- | --- |
| Kernel | Susc | Infe | Model | reliability (%) | mean bias | mean post-prior | mean post-prior |
| wide | broadleaf | broadleaf | homog | 95 | 0.04 |  |  |
|  |  |  | heter | 95 | 0.02 | +0.29 | --0.05 |
| wide | broadleaf | arable | homog | 100 | 0.09 |  |  |
|  |  |  | heter | 100 | 0.02 | +0.37 | -0.03 |
| moderate | broadleaf | broadleaf | homog | 90 | 0.07 |  |  |
|  |  |  | heter | 100 | 0.02 | +0.43 | +0.01 |
| moderate | broadleaf | arable | homog | 70 | 0.10 |  |  |
|  |  |  | heter | 80 | 0.04 | +0.45 | +0.07 |

In the ‘entire outbreak’ data availability setting, it is no longer meaningful to consider predictive performance on held-out future outbreak data. Parameter recovery is assessed for the same set of parameter combinations as for the ‘early stages’ setting. Reliability of inference for the transmission kernel parameter is excellent for the heterogenous model but less good for the homogenous model (Table 3). This parameter is consistently overestimated but especially by the homogeneous model, which again is consistent with what was observed in (A. Cook et al. 2007). The posterior medians of the (large) susceptibility and infectivity parameters have shifted closer to their true values when compared to Table 2. This is as we would expect given that we have more data available to fit the model in this setting. In particular, there is now clearer evidence of a shift in the infectivity parameter posterior distribution away from the prior distribution.

**Table 3.** Full outbreak parameter recovery. Models are fitted to data on entire simulated outbreaks on the Estonian landscape. For other details, see text for Table 2. Mean post-prior is the mean, over replicates of the posterior median minus the prior median for the parameter.

| Scenario |  |  |  | Kernel parameter |  | Susceptibility | Infectivity |
| --- | --- | --- | --- | --- | --- | --- | --- |
| Kernel | Susc | Infe | Model | reliability (%) | mean bias | mean post-prior | mean post-prior |
| wide | broadleaf | broadleaf | homog | 75 | 0.19 |  |  |
|  |  |  | heter | 95 | 0.05 | +0.64 | +0.05 |
| wide | broadleaf | arable | homog | 65 | 0.20 |  |  |
|  |  |  | heter | 95 | 0.08 | +0.58 | +0.05 |
| moderate | broadleaf | broadleaf | homog | 75 | 0.13 |  |  |
|  |  |  | heter | 90 | 0.03 | +0.63 | +0.12 |
| moderate | broadleaf | arable | homog | 70 | 0.16 |  |  |
|  |  |  | heter | 95 | 0.04 | +0.61 | +0.15 |

Finally, to put these results into context, we take a closer look at the spatial variation in the Estonian landscape, which we have used for these simulations. At the 10km resolution, the Estonian landscape is not particularly heterogeneous in terms of land use (Figure 4). Most of the 575 10km square patches comprise a mixture of land uses, with very few patches consisting mostly of a single land use. The low heterogeneity in land use is one factor leading to limited variation in estimated patch susceptibilities (Figure 5). In general, we might expect that increased heterogeneity in land use, increased differences in susceptibility/infectivity between land uses, a wide kernel and higher numbers of available case reports are all factors that should increase the benefits of modelling spatial variation in susceptibility and infectivity.

**Figure 4.**
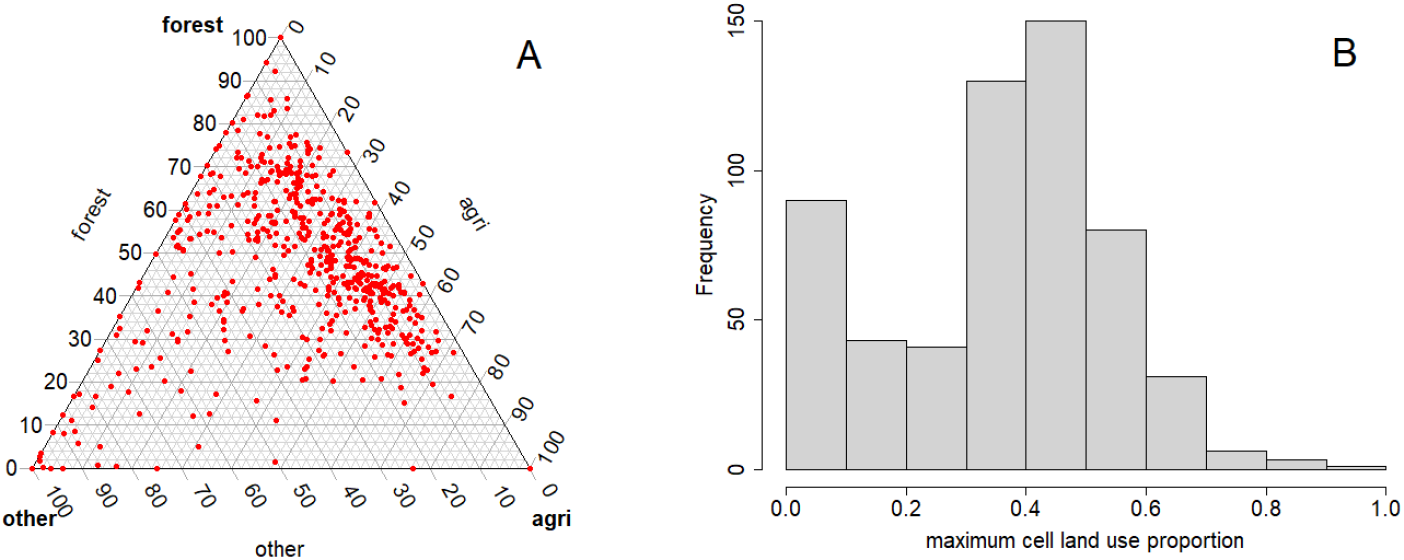
Landscape heterogeneity in Estonia. Left: ternary diagram showing the land use composition of every 10km square patch covering Estonia (land use amalgamated into three categories – forest, agricultural and other – for ease of visualisation). Right: histogram showing maximum patch land use proportion for every 10km square patch in Estonia, with six land uses as in the model covariates.

**Figure 5.**
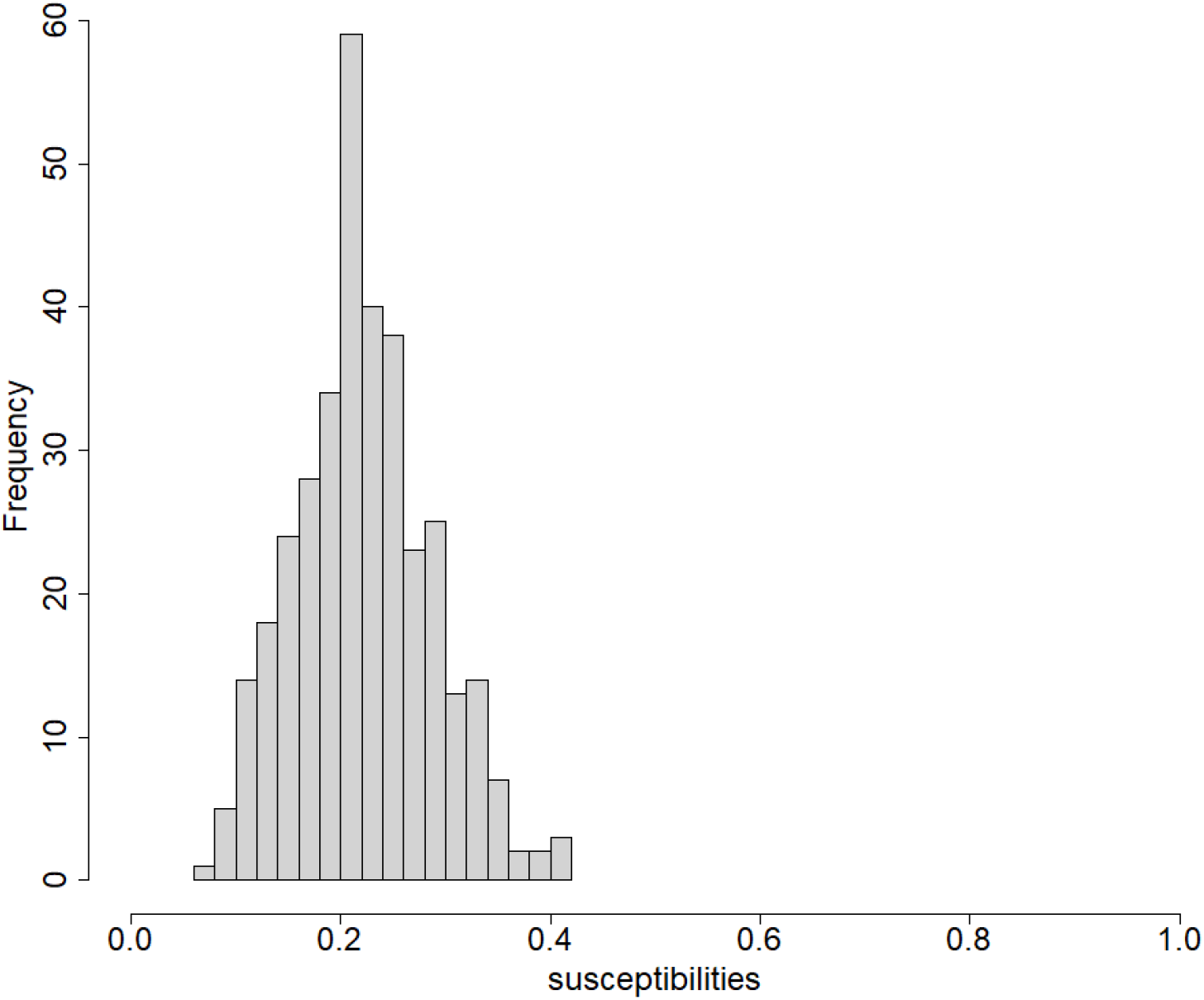
Typical distribution of (inferred) posterior mean patch susceptibilities given data from a simulated disease outbreak in Estonia. There is some variation but few patches with very high or low susceptibilities.

## 4. Case study: African Swine Fever in the Baltic states

### 4.1 Background/approach

Here, we used all ASF cases in wild boar populations reported in Estonia between 2014 and 2019 to the World Organisation for Animal Health (WOAH). The Estonian ASF outbreak had very serious impacts for both the pig industry and wildlife. African Swine Fever leads to high mortality, and its spread is difficult to control. It continues to pose a major ongoing threat in many regions of Europe (Cwynar, Stojkov, and Wlazlak 2019; Schulz et al. 2019; Depner et al. 2017). In this study we focus on better quantifying risk of transmission and spread of ASF in the Estonia wild boar population. Knowledge of the wild boar population in this region is incomplete, though estimated wild boar density maps have been published (Depner et al. 2017). Within Estonia, there have been extensive efforts to control the ASF outbreak, partly by reducing the density of wild boar. These efforts are reflected in density maps showing an apparent reduction in the population over time (Depner et al. 2017). Case reports of ASF in dead wild boar are available at monthly time intervals, with detailed location information. Therefore, given the uncertainty in knowledge of the Estonian wild boar population described above, we apply the iPAR framework to this outbreak.

To apply the framework to the ASF outbreak in Estonian wild boar, we must first define a grid of patches covering Estonia, as well as covariates likely to be related to wild boar density and hence to patch susceptibility and infectivity. Land use is one of the key factors likely to determine wild boar density (Tarvydas and Belova 2022), and hence we use high resolution CORINE land use data available for Estonia (CLMS 2018). The case report data for wild boar in Estonia also have high spatial resolution. However, if the patches used to define the infectious units of the model are too small then (a) fitting the model will be very computationally intensive, and (b) the persistence assumption inherent in our assumed SI framework is less likely to be reasonable. As a compromise we initially opted for square patches of size 10km by 10km, though smaller patches would also be feasible. This choice of patch size was informed by a persistence of infection analysis, which revealed that, at the 10km resolution, ASF in wild boar in Estonia typically persists for long periods of time, often multiple years (ESM Appendix 10). Any of the 10km patches composed wholly of sea or inland water were removed from the set of patches. In the case of Estonia, there are 575 patches covering the country. Figure 6 illustrates the spatial spread of ASF through the wild boar population in Estonia at the 10km patch resolution.

**Figure 6.**
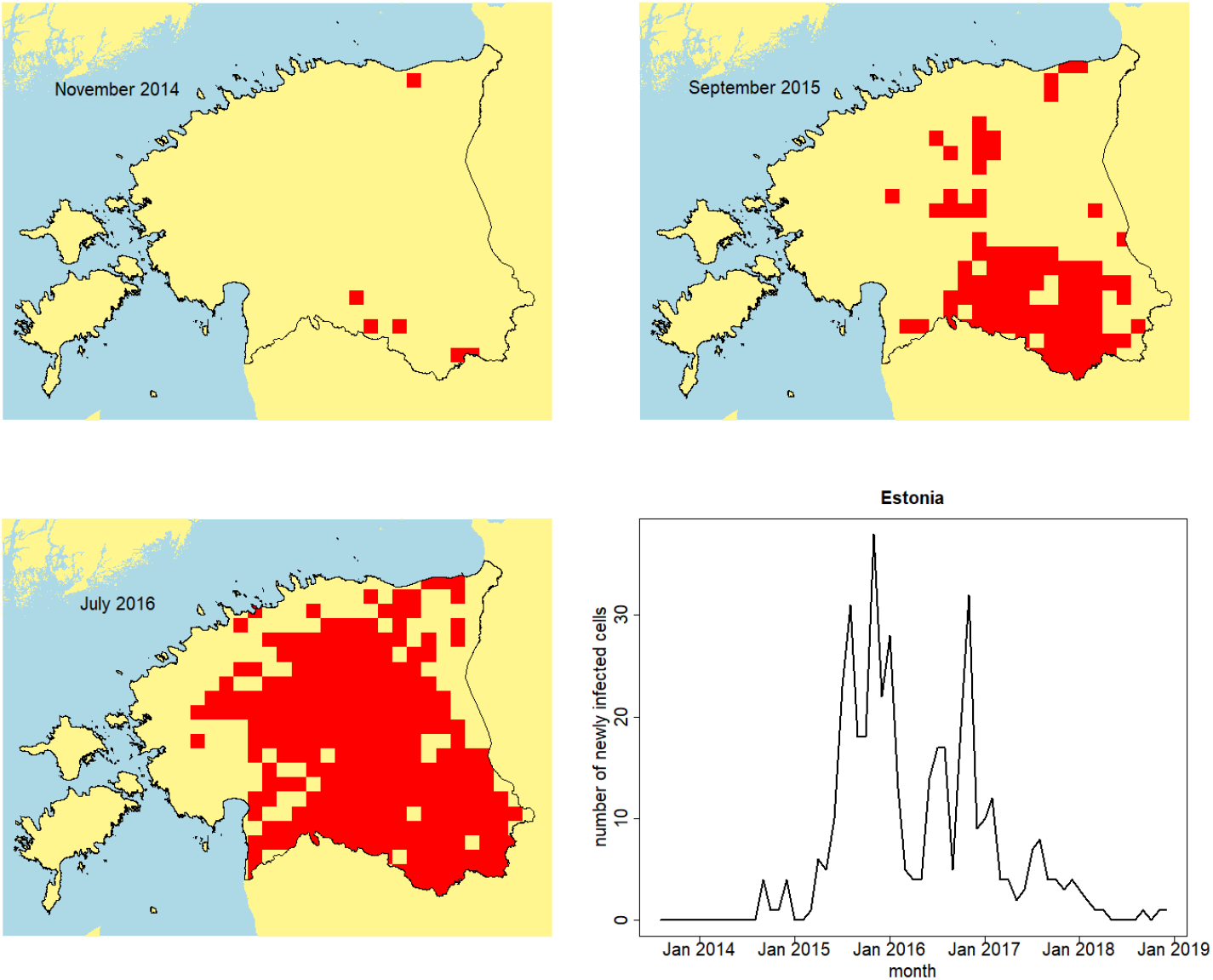
Case reports: Spatially aggregated African Swine Fever wild boar cases reported in Estonia. Snapshots at: November 2014, September 2015, July 2016. Bottom left panel shows the outbreak incidence curve, generated using spatially aggregated wild boar case reports.

We took CORINE land cover data (CLMS 2018) at 100 metre resolution and spatially aggregated it to the 10km patches to obtain proportions of each land use category for each patch. We also aggregated some of the CORINE land cover categories, resulting in a total of 6 categories: 1=urban, 2=agriculture, 3=broadleaf/mixed forest, 4=coniferous forest, 5=semi-natural, 6=wetlands/water bodies excluding sea. We defined the susceptibility of a patch *i* to be 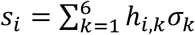 where the *σ*_*k*_ are parameters to be estimated and *h*_*i,k*_ is the proportion of land in patch *i* occupied by land use category *k*. Similarly, consistent with the notation in Section 2.1, the infectivity of a patch *i* is defined as 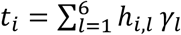.

A case report from the outbreak is associated with a specific month. So, the time of the case report is considered to be interval-censored, with the time interval equal to the relevant month. The model can now be fitted to the outbreak data as described in Section 2.2.1. We could have taken into account other covariates but, for simplicity, we chose to use land use data only. This case study is intended as a proof of principle, rather than a definitive analysis of ASF transmission in wild boar in Estonia.

### 4.2 Parameter estimates

The constant-in-time and varying-in-time heterogeneous models were fitted to the full set of Estonian outbreak data, with the estimated parameters as in Table 4. The *DIC*_1_ metric was computed for both model fits, and was found to strongly favour the varying-in-time model (details in ESM Appendix 4). This is not surprising given the strong decreasing trend in *h*(*t*) that was identified (Table 4). Generally, parameter estimates were consistent between the two model variants. There is much uncertainty, but broadleaf woodland seems to have the highest susceptibility, while agricultural land has the highest infectivity. External transmission occurs at very low levels, and the estimated kernel is relatively narrow. The estimates of the overall rate *ρ* for the two models are not easily comparable because, in the varying-in-time model, *ρ* is multiplied by the function *h*(*t*) which is estimated as less than one for most of the observation time interval. So it is not surprising that the estimated value of *ρ* is higher for the varying-in-time model.

### 4.3 Model predictions

Finally, we assess the predictive performance of the model fitted to the Estonian data. We fit the constant-in-time model to the early stages of the outbreak in Estonia, holding out data from later stages of the outbreak in order to test the model’s predictions. Both the heterogeneous and homogeneous variants of the model were fitted.

The predicted number of infected patches under the heterogeneous and homogeneous models tends to be overestimated, though the predictions of the heterogeneous model lie a little closer to the actual data (Figure 7A). The predicted number of infected patches from simulation of a heterogeneous varying-in-time model fitted to the entire outbreak is much closer to the actual data (Figure 7B). This indicates that the varying-in-time model is sufficiently flexible to capture the key trends in the observed cumulative incidence curve in Estonia.

**Figure 7.**
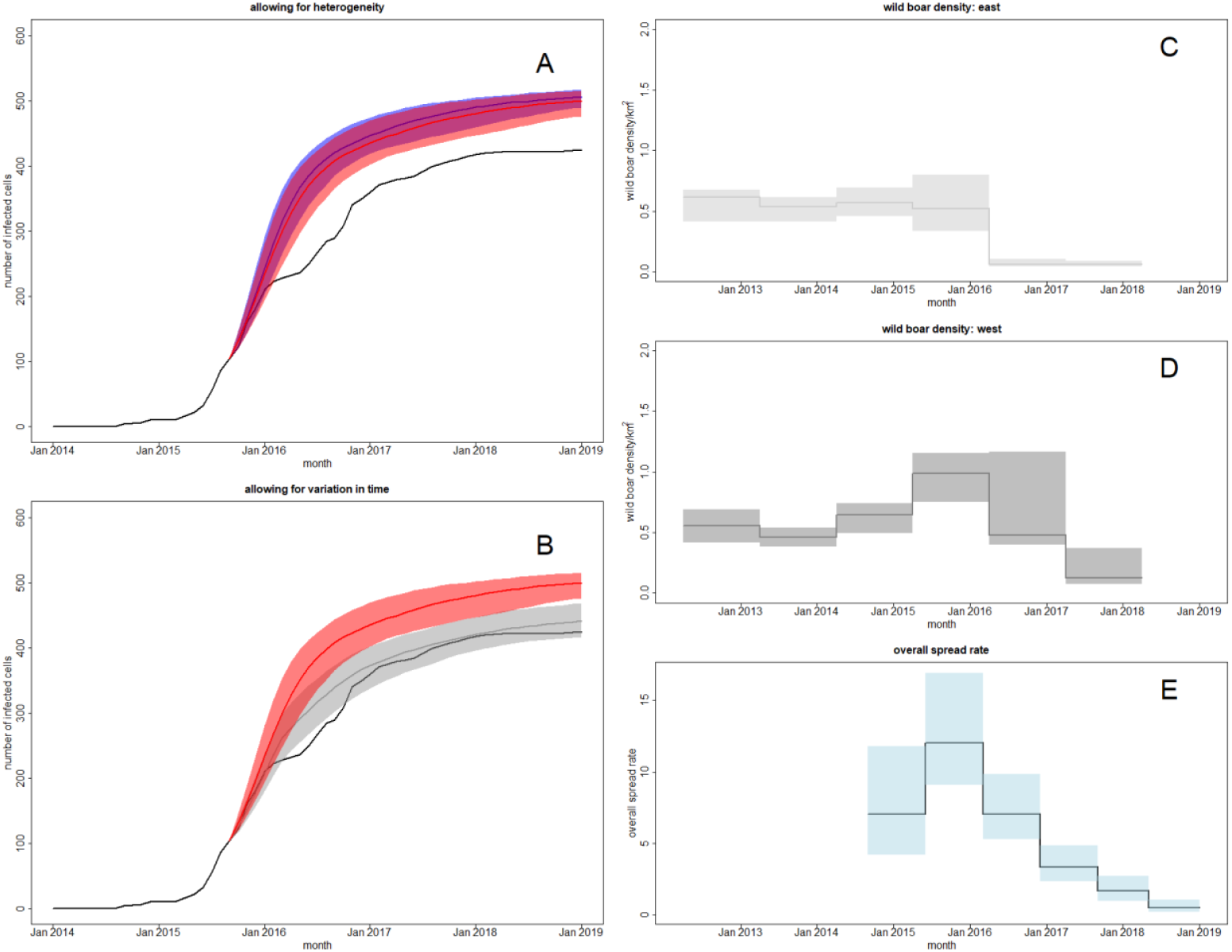
Model prediction and validation. *Left column*: Predicted numbers of infected 10km patches from models fitted to the early stages of the ASF outbreak in Estonian wild boar. Two constant-in-time models (heterogenous, homogeneous) were fitted to outbreak data up to September 2015. The black lines above show the true cumulative incidence curve at 10km patch resolution. A: the blue line and region show the posterior predictive median and 95% credible interval for the total number of infected patches under the homogeneous model. The red line and region show the corresponding quantities under the heterogenous model. B: the red line and region show the posterior predictive median and 95% credible interval for the total number of infected patches under the heterogeneous model. The grey line and region show the corresponding quantities under a varying-in-time heterogeneous model fitted to the full set of outbreak data. *Right column*: External validation via hunting bag data from (Schulz et al. 2019). C: estimated wild boar density in the east of Estonia. D: estimated wild boar density in the west of Estonia. E: estimated overall spread rate *ρh*(*t*) from the varying-in-time model fitted to the entire ASF outbreak in Estonia. The black line represents the posterior median, the blue area shows the 95% credible interval.

### 4.4 External validation

Many aspects of the fitted model are difficult to validate externally. However, one aspect of interest is the time varying function *h*(*t*), which was estimated to initially increase from the baseline value of 1 in 2015 to a maximum of 1.71 in 2016, before dropping to 0.06 by the end of the outbreak (Table 4). The function is intended to model the effects of disease control measures, and in Estonia a key control measure was reduction of the wild boar population. The trend seen in the estimated time varying function is broadly consistent with reported trends in wild boar population density across Estonia based on information from hunters (see Figure 5 in (Schulz et al. 2019)). This comparison provides some evidence for the validity of the fitted varying-in-time model (see Figures 7C-E).

### 4.5 Quantitative risk assessment for disease control

In addition to the retrospective analysis described above which shows the impact of the control measures that were adopted, the iPAR framework also provides information relevant to the within-outbreak targeting of disease control activities. For example, Figure 8A illustrates the estimated susceptibilities of patches in Estonia for the constant-in-time model, identifying which locations are more likely to be susceptible to disease incursion. Although there are some areas with greater susceptibilities there does not seem to be a strong trend in susceptibility across the country. However, Table 4 also provides potentially useful information to inform disease control efforts by identifying the land use classes (broadleaved and agricultural) that contribute most strongly to susceptibility and infectivity. Such information may be useful in guiding development of control policies and programmes.

**Figure 8.**
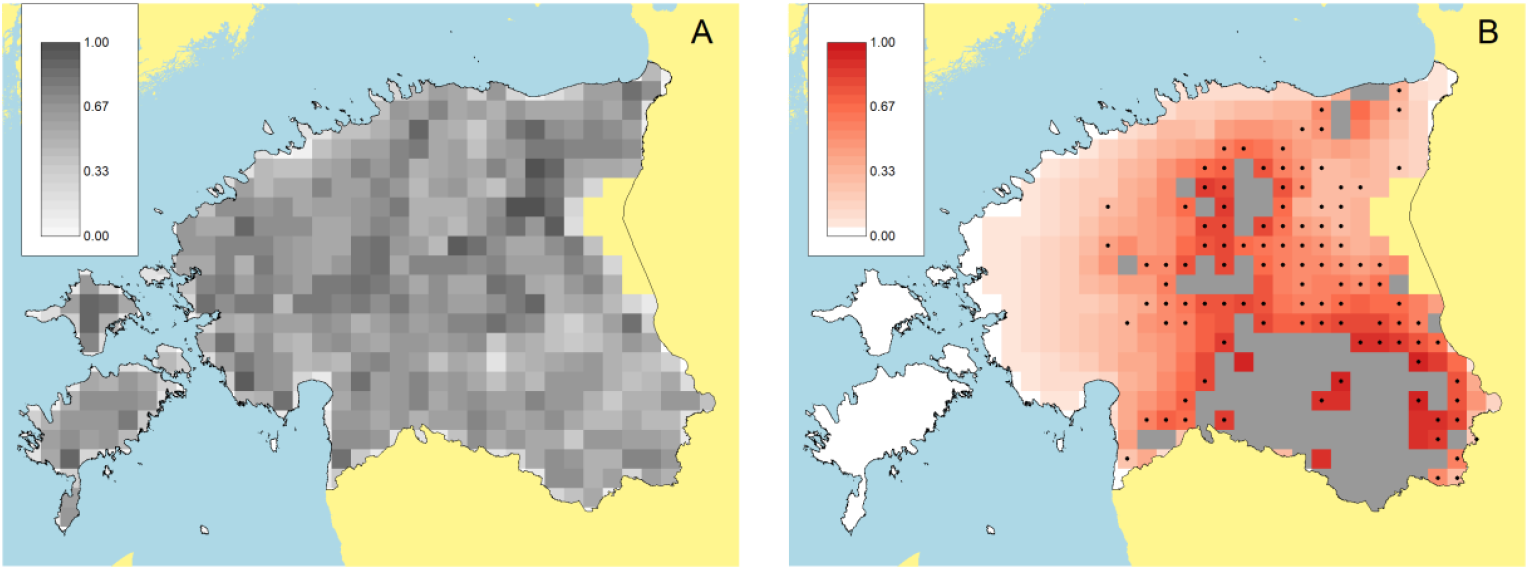
Quantifying disease risk. (A) Susceptibility map for Estonia obtained from fitting the constant-in-time model to the Estonian ASF wild boar outbreak data. (B) Risk map illustrating predictions from the constant-in-time heterogeneous model fitted to the early stages of ASF outbreak in wild boar in Estonia. Grey patches are recorded as infected at the end of the early stage of the outbreak. Red colours indicate probability of infection four months beyond this end point. The black dots show patches that did in fact become infected within this time horizon. Alternative summaries of the same predictions may be found in Table 5 and Figure 7 (left).

In addition, risk maps can be helpful for visualising spatial predictions of which areas will become infected over defined scenarios and specified time horizons. Based on observation of the early stage epidemic, up to September 2015 (grey patches), Figure 8B shows model predictions of the probability of future infection. Such information can aid decisions about where to focus limited resources for disease control as the outbreak unfolds in real time. This prediction is over a specified time horizon under the assumption that rates of spread remain fixed. However, we also note that such use of this information means that actual spread will ideally be different from that predicted. This is due to the impact of disease controls as illustrated in FigureS 7C-E which suggests a substantial benefit of disease control onwards from 2016.

**Table 4.**
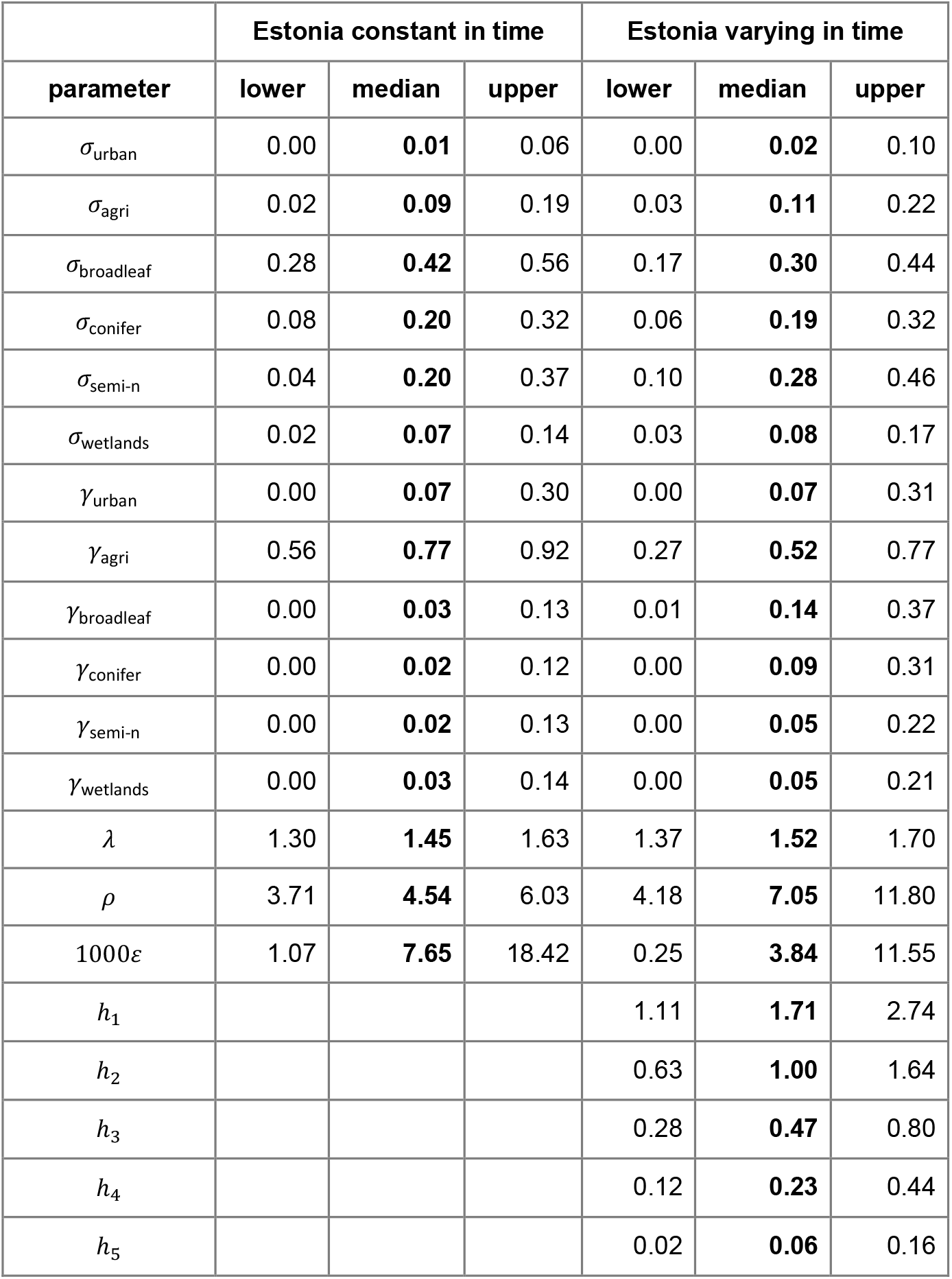
Parameter estimates obtained from fitting the iPAR model to data from the entire Estonian outbreak: posterior median and upper/lower limits of 95% credible intervals. The parameters *h*_0_ = 1, *h*_1_, …, *h*_5_ represent the piecewise-constant time-varying function *h*(*t*) as described in Section 2.2.1. The time intervals over which the function is constant are pre-specified and are as depicted in Figure 7 (bottom right panel).

|  | Estonia constant in time |  |  | Estonia varying in time |  |  |
| --- | --- | --- | --- | --- | --- | --- |
| parameter | lower | median | upper | lower | median | upper |
| $\sigma_{\text{urban}}$ | 0.00 | **0.01** | 0.06 | 0.00 | **0.02** | 0.10 |
| $\sigma_{\text{agri}}$ | 0.02 | **0.09** | 0.19 | 0.03 | **0.11** | 0.22 |
| $\sigma_{\text{broadleaf}}$ | 0.28 | **0.42** | 0.56 | 0.17 | **0.30** | 0.44 |
| $\sigma_{\text{conifer}}$ | 0.08 | **0.20** | 0.32 | 0.06 | **0.19** | 0.32 |
| $\sigma_{\text{semi-n}}$ | 0.04 | **0.20** | 0.37 | 0.10 | **0.28** | 0.46 |
| $\sigma_{\text{wetlands}}$ | 0.02 | **0.07** | 0.14 | 0.03 | **0.08** | 0.17 |
| $\gamma_{\text{urban}}$ | 0.00 | **0.07** | 0.30 | 0.00 | **0.07** | 0.31 |
| $\gamma_{\text{agri}}$ | 0.56 | **0.77** | 0.92 | 0.27 | **0.52** | 0.77 |
| $\gamma_{\text{broadleaf}}$ | 0.00 | **0.03** | 0.13 | 0.01 | **0.14** | 0.37 |
| $\gamma_{\text{conifer}}$ | 0.00 | **0.02** | 0.12 | 0.00 | **0.09** | 0.31 |
| $\gamma_{\text{semi-n}}$ | 0.00 | **0.02** | 0.13 | 0.00 | **0.05** | 0.22 |
| $\gamma_{\text{wetlands}}$ | 0.00 | **0.03** | 0.14 | 0.00 | **0.05** | 0.21 |
| $\lambda$ | 1.30 | **1.45** | 1.63 | 1.37 | **1.52** | 1.70 |
| $\rho$ | 3.71 | **4.54** | 6.03 | 4.18 | **7.05** | 11.80 |
| $1000\varepsilon$ | 1.07 | **7.65** | 18.42 | 0.25 | **3.84** | 11.55 |
| $h_1$ | | | | 1.11 | **1.71** | 2.74 |
| $h_2$ | | | | 0.63 | **1.00** | 1.64 |
| $h_3$ | | | | 0.28 | **0.47** | 0.80 |
| $h_4$ | | | | 0.12 | **0.23** | 0.44 |
| $h_5$ | | | | 0.02 | **0.06** | 0.16 |

**Table 5.** Predictive performance of the constant-in-time model fitted to data from the early stages of the ASF outbreak in wild boar in Estonia. For each fitted model (homogeneous/heterogeneous) the table shows AUC (area under the ROC curve) and the true positive rate at 5% false positive rate (TPR @ 5% FPR). All metrics were evaluated at a time of four months beyond the last observations used for fitting the models.

| Model | Discrimination metrics |  | Spatial extent of infection |  |
| --- | --- | --- | --- | --- |
|  | AUC | TPR @ 5% FPR | 2.5%ile | 97.5%ile |
| homog | 0.90 | 0.44 | 0.97 | 1.39 |
| heter | 0.90 | 0.47 | 0.92 | 1.34 |

By comparing spatial risk prediction to actual case reports over a limited time horizon we see the heterogeneous model identifies future infected patches slightly better than the homogeneous model (Table 5). However, the differences between the two models’ predictions are not substantial. One reason for this observation is likely to be that the landscape of Estonia is relatively homogeneous in the sense that there is little spatial variation in susceptibility at large scales (see Section 3.4). Another explanation may be the narrowness of the estimated kernel, which is much narrower than the kernels used in the simulation experiments in Section 3.4. Further simulations (ESM Appendix 11) similar to those in Section 3.4 but with a narrow kernel give predictive performance similar to that in Table 5.

## 5. Discussion

This paper develops the novel iPAR (inference for populations at risk) framework for inferring and modelling spread of disease in an unknown population at risk, given outbreak data in the form of only case reports with possibly less than precise spatial and temporal coordinates. The method introduced exploits the intuitive idea that case reports simultaneously provide information on both disease spread and the population at risk, enabling modelling in relatively data poor scenarios not previously amenable to inference based on mechanistic transmission models. Where it is available the iPAR framework also allows for the use of spatial covariate data that may be informative about the population at risk. iPAR uses a patch-based model with patch susceptibility and infectivity varying across the region over which disease is spreading. By focussing on estimation of patch susceptibilities and infectivities we circumvent any requirement for detailed knowledge of the spatial distribution of the population at risk. The use of patches also means that less than precise location information is required and Bayesian inference via data-augmented MCMC enables estimation of uncertainty in infection times associated with time censoring of case reports.

We demonstrate that the inference techniques employed here reliably estimate model parameters, with uncertainty reflecting available case data. Typically parameters in the transmission kernel, as well as external transmission rates, are well estimated, susceptibility parameters are moderately well estimated, while as is to be expected infectivity parameters are the least well estimated (see e.g. Pooley et al. 2020). The iPAR approach is also able to estimate the ‘overall’ rate of infection *ρ*, and time variation in this quantity captured by *ρh*(*t*). In the case study this estimated time variation was shown to correlate with impacts disease control efforts to reduce wild boar density in Estonia. Application of the iPAR approach is thus able to retrospectively characterise an outbreak and assess the likely benefits of control actions taken. However, perhaps more critically iPAR can also be used to provide information useful for risk assessment during an outbreak. In the ASF case study presented here this includes: (i) identifying which land use classes and thus areas pose the highest intrinsic disease risks – potentially enabling targeted controls; (ii) quantifying the scale of spatial spread – information that could be used to design protocols for exclusions zones, ring vaccination or culling; and (iii) spatio-temporal quantification of future risk – identifying when and where new cases are most likely to arise thereby directing effort to where it is most needed in real time.

A key aspect of the iPAR framework is the ability to account for variation in susceptibility and infectivity across the landscape. Simulations highlighted scenarios where the heterogeneous model, with varying susceptibility/infectivity, significantly outperforms the corresponding homogeneous model. However, in our case study the heterogeneous model offered only a minimal improvement over the homogeneous model, but was nonetheless favoured by model selection. It is possible that use of other covariates not available to our analysis, might give a heterogeneous model that offers more of an improvement, but the narrowness of the estimated kernel makes this less likely. Intuitively, with a narrow kernel, there are fewer possible destinations for onward transmission of disease, so having a heterogeneous model that attempts to distinguish these possible destinations becomes less useful.

The most appropriate spatial resolution at which to model the spread of disease is dependent on multiple factors. If the spatial resolution is too low then the model may not be able to produce accurate estimates of quantities of interest (Mills and Riley 2014), but if the resolution is too high then there may be computational issues and a lack of suitable outbreak/covariate data. Other factors to consider are the intended application, model assumptions and the key quantities of interest. In the case study presented here, the *SI* model assumes persistence of infection, which is less likely to be reasonable at very high resolutions. We chose 10km resolution for the case study as analysis demonstrated good persistence at this scale. Furthermore preliminary analysis of the case study data at 5km resolution gave similar results (see ESM Appendix 9).

To handle model parameters constrained to the simplex, e.g. patch land use susceptibilities, this work made novel, within the context of transmission models, use of methods from compositional data analysis, such as the isometric logratio transform. As noted previously, there is typically slight overestimation of the ‘overall’ rate of infection *ρ*, which is proportional to the rate of transmission between two uniform patches (patches containing equal proportions of all land uses). Investigation of this phenomena suggests that it arises from the influence of the prior distribution on parameters for which the available data are not strongly informative (see ESM Appendix 7). However, we note here that the properties of transmission models containing such constrained parameters do not seem to have been well explored in the epidemic modelling literature. Issues for further research include specification of prior distributions for parameters on the simplex, and interpretation of credible intervals for parameter components that are highly constrained and interdependent. In scenarios where knowledge of the population at risk is limited, compositional covariates such as land use proportions may be the only data that we have available, so methods that make best use of such data could have significant practical impact on disease control in data poor scenarios such as those addressed here.

Future methodological developments could extend the implementation of the iPAR framework to include more general disease dynamics beyond the susceptible-infected dynamics employed here. In addition, it would be desirable to implement iPAR for multiple host species e.g. transmission of ASF in wild boar and domestic pigs, or transmission of *Xylella fastidiosa* in multiple susceptible plant species (?). Nonetheless there are numerous potential applications of the implementation of the iPAR framework introduced here. These include not only the study of other outbreaks of ASF in wild boar and/or domesticated pigs, but also other transmissible disease outbreaks where case only data are available. These might include Newcastle disease in poultry or highly pathogenic avian influenza in wild birds and/or backyard poultry.

In conclusion, we have developed a modelling framework that fills a gap in epidemiological modelling approaches. The iPAR approach allows modelling of spatial disease spread when the distribution of the disease host is uncertain, and the model can be parameterised using only disease case report data. In recent years, the amount of data available to inform models for spatial disease spread has rapidly increased. At the same time, data availability and data privacy continue to be an issue in many situations. Our approach can circumvent the requirement for host distribution data and so can open up many disease outbreak datasets for analysis.

## Code

Computer code for performing the analyses described in this manuscript may be found at https://github.com/spcatterall/iPAR.

## Appendix 1

### Sampling algorithm

Here we provide further details of the MCMC algorithm used to obtain samples from the joint posterior distribution *P*(*θ*, 𝒰|𝒯) of the model parameters *θ* and the actual unobserved patch infection times 𝒰, where 𝒯 denotes the set of interval-censored patch infection times which we assume are provided to us as data. The algorithm initialises and simulates a Markov chain whose stationary distribution is the posterior distribution. At each iteration of the algorithm, updates are proposed to each parameter and infection time in turn. A more precise description follows, in which the superscript (*m*) indicates the value of a quantity at the *m*’th iteration of the algorithm. We use the same notation as used in Section 2 of the manuscript.

#### Initialisation

Initialise parameters with values that are arbitrary but also consistent with the selected prior distributions. For a patch *i*, let *T*_*i*_*ϵ*{0,1,2, …, *N* + 1} denote the ‘time category’ into which the patch falls. Then, for 1 ≤ *T*_*i*_ ≤ *N*, we initialise the infection time of the patch uniformly at random on the time interval (*t*_*j*−1_, *t*_*j*_]. If *T*_*i*_ = 0 then the patch is taken to be infected at the beginning of the modelled time period. If *T*_*i*_ = *N* + 1 then the patch is taken to be uninfected at the end of the modelled time period.

#### Iteration

Repeat the following sequence a large number of times:

a. Evaluate the current likelihood. In the manuscript this is denoted *P*(𝒯, 𝒰|*θ*) but for simplicity we denote it here by *L*. Propose a new value for the transmission parameter *λ* by sampling from the proposal distribution *q*(*λ*^(*m*)^, *λ*^*^). Specifically, we take 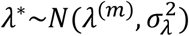 which is a normal distribution centred on the current value of the parameter. Recalculate the likelihood with the proposed parameter value, to give a new likelihood *L*^*^. The proposed change is accepted with Metropolis-Hastings acceptance probability

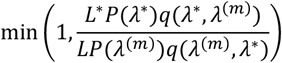 In this case, we take *λ*^(*m*+1)^ = *λ*^*^ and the likelihood is updated to its new value *L*^*^. If the change is rejected then we take *λ*^(*m*+1)^ = *λ*^(*m*)^ and the likelihood remains the same. Note that, because the proposal distribution is symmetric, the two proposal factors in the acceptance probability cancel, which simplifies the calculations.
b. Repeat (a) for the other model parameters in turn: overall rate *ρ*, background rate *ε*, time varying parameters *h*_*i*_ (for the varying-in-time model), the susceptibility parameters 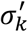 and the infectivity parameters 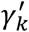.
c. Sample the simplex parameters *σ* and then *γ*. The components of *σ* are constrained to sum to 1, so we can’t sample the components independently in the same way that, for example, the susceptibility parameters 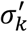 can be sampled independently, one at a time. Instead, we use a joint proposal distribution *q*(*σ*^(*m*)^, *σ*^*^). Specifically, we take

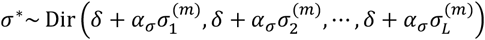 This is a Dirichlet distribution with a specified set of concentration parameters. It is the analogue, for parameters on the simplex, of the Gaussian random walk proposal used in (a) for parameters defined on real intervals. The acceptance probability and other aspects of the procedure are the same as in step (a). However, in this case the proposal distribution is not symmetric, so the two proposal factors in the acceptance probability do not cancel. The sampling process is then repeated for the other simplex parameter *γ*.
d. Sample the unobserved patch infection times *U*_*i*_, taking each one in turn. This is carried out in the same way as in (a), with a normal proposal density 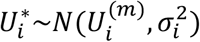 and acceptance probability

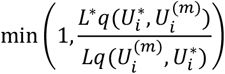

#### Optimisation

The first 10000 iterations of each Markov chain are discarded as ‘burnin’. During the burnin phase, the proposal distribution parameters such as 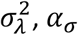 and 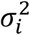 are adapted to ensure that the acceptance probabilities are not too large or small. This is done in a similar manner to that described in Pooley et al. (2020). The parameter *δ* in the Dirichlet proposal was fixed at 1, a value obtained by trial and error; it was not adapted in the same way as the other proposal distribution parameters. Its purpose is to stabilise the algorithm, in particular to prevent proposed values from getting too close to the boundary of the simplex.

When the likelihood is recalculated during the MCMC, only the parts of the likelihood that might actually change are computed. This ensures that the computation time is kept to a minimum.

#### MCMC convergence

Typically, five chains are run in order to fit the model. The Gelman-Rubin convergence test is used to assess convergence, along with inspection of trace plots, autocorrelation, between-parameter correlation and estimation of the effective sample size.

## Appendix 2

### Kernel normalisation

The transmission kernel is a truncated power law *K*(*d*) = *d*^−2*λ*^*I*(*d* < *d*_max_)/*m*(*λ*) where *m*(*λ*) is a normalisation constant defined as follows.

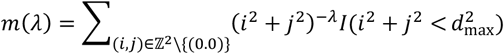

Intuitively, *m*(*λ*) is the sum of the force of infection described by the unnormalized kernel, given a single source of infection at coordinates (0,0) and susceptible units arranged on a two-dimensional integer lattice. If we divide the kernel by this sum, it means that changing *λ* will only change the spatial distribution of the force of infection arising from a single infectious unit, rather than changing the total force of infection on the surrounding landscape. This leads to better separation of the effect of *λ* from the effect of the overall rate of infection *ρ*.

## Appendix 3

### Prior distributions

The prior distributions for the model parameters were taken to be uninformative, and were generally chosen to be uniform distributions over their chosen supports. Most parameters had independent prior distributions, with the exception of the two simplex parameters where it was essential to allow for dependence between the simplex components. To fit the iPAR model, whether time-varying or constant-in-time, to the Estonian data in the case study, we chose the following prior distributions.

| Parameter | Prior distribution |
| --- | --- |
| $\sigma, \gamma$ | Uniform on the simplex $S^6$ |
| $\lambda$ | Uniform on the interval (0,10) |
| $\rho$ | Uniform on the interval (0,10000) |
| $\varepsilon$ | Uniform on the interval (0,10) |
| $h_i$ | Uniform on the interval (0.1,100) |

## Appendix 4

### DIC computation and values

In this appendix we use the same notation as in Gamado et al. (2017). In that paper, the DIC metric *DIC*_1_ is defined by the following formula

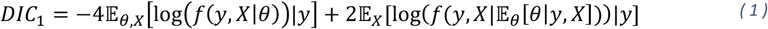

Here, *f* is the complete/full likelihood, *y* represents the observed data, *X* represents the unobserved data, while *θ* is the model parameters. This metric corresponds to *DIC*_4_ in Celeux et al. (2006). For each value of *X*, the second term requires evaluation of a posterior expectation (conditional on that value of *X*), so *DIC*_1_ is quite expensive to compute if there is no closed form available, or if the dimension of *θ* is high. However, inspection of the supplementary material of Gamado et al. (2017) shows that, in fact, the following metric was computed

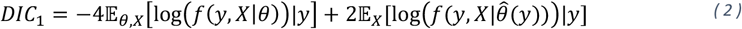

Here 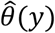 is the posterior expectation of the parameter vector given the data. This metric is a version of *DIC*_6_ in Celeux et al. (2006). It is this latter metric that has been computed for the fitted models in the manuscript, but with 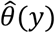 being the posterior median rather than expectation. When we refer to *DIC*_1_ in the manuscript, it is this second definition that we are assuming.

The computation of *DIC*_1_ is an additional task which can be carried out after the MCMC algorithm has been completed. During the MCMC we store the loglikelihood values obtained at each iteration. Averaging these stored loglikelihood values gives the first expectation in (2) above. The second expectation requires a bit more work. First, we compute the posterior medians 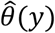. Then, at each iteration/sample we compute the loglikelihood evaluated at 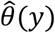 and the ‘current’ set of latent variables. Averaging these loglikelihoods over all iterations gives the second expectation in (2). Note that the samples can be thinned before carrying out these computations, which substantially reduces computation time.

#### *DIC*_1_ values for Section 3.3

More context regarding the simulated scenarios listed below is provided in the main text. The lowest *DIC*_1_ for each scenario, which is the *DIC*_1_ of the favoured model, is highlighted in bold.

| Scenario | Constant-in-time model | Varying-in-time model |
| --- | --- | --- |
| 1 (linear decrease) | 2908 | **2874** |
| 2 (constant) | **2882** | 2886 |
| 3 (sudden drop off) | 2961 | **2935** |

#### *DIC*_1_ values for Section 4.2

More context regarding the data used to fit these models is provided in Section 4, the case study. The lowest *DIC*_1_, which is the *DIC*_1_ of the favoured model, is highlighted in bold.

| Constant-in-time model | Varying-in-time model |
| --- | --- |
| 3203 | **2987** |

## Appendix 5

### the simplex and the isometric logratio transformation

The disease transmission model described in the manuscript uses parameters for susceptibility and infectivity that are constrained to the simplex 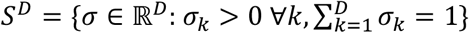, where *D* = 6 in the case study. Estimating a parameter constrained to the simplex can be challenging due to counterintuitive effects. Two possible effects are the following:

1. It can be difficult to estimate parameter values that are on or close to the boundary of parameter space, and credible intervals as commonly constructed may not contain the true parameter value. So, we can expect difficulties if a simplex parameter is close to the boundary of the simplex. One approach that can sometimes help is the ilr transform, which we discuss below.
2. The prior distribution on the simplex is assumed to be uniform, which implies that the prior mean is the centroid of the simplex. If the data are not very informative then simplex parameter estimates are likely to be drawn towards the centroid somewhat.

Below, we consider parameter estimation for a simple multivariate normal model on the simplex. Our purpose here is to highlight that the above two effects can be relevant even for very simple models. Then, we consider the above two effects for the disease transmission model used in this manuscript.

#### The isometric logratio transform

The isometric logratio transform ilr: *S*^*D*^ → ℝ^*D*−1^ is defined by

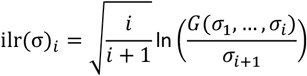

where *G* denotes the geometric mean and 1 ≤ *i* ≤ *D* − 1. This transform is helpful because it maps simplex parameters to more familiar real vector-valued parameters. On the other hand, the components of ilr(σ) may not be easy to interpret in terms of the components of σ.

#### The multivariate normal simplex model

The model is as follows. It has a single parameter *μ* ∈ *S*^*D*^. Observed data are assumed to be independent samples from the normal on *S*^*D*^ distribution 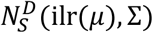, which was introduced by Mateu-Figueras et al. (2013). This distribution is essentially a multivariate normal distribution on the simplex and has density

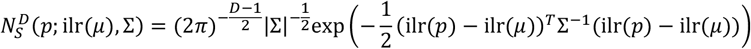

Where, for simplicity, the covariance matrix Σ is assumed known and equal to *σ*^2^*I*. In summary, the observed data points are 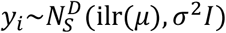 for 1 ≤ *i* ≤ *n*.

Suppose also that we assume a spherically symmetric prior for *μ* of the form 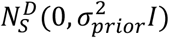. The posterior distribution for *μ* can then be computed analytically in the same way as for multivariate normal data (see e.g. Gelman et al. 2003) to obtain 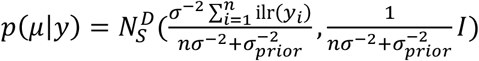. The mean vector is a convex combination of the prior mean in real space (zero) and 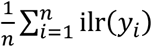, as expected. Thus, even if 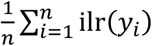 is close to ilr(*μ*), the mean of the posterior will be drawn somewhat towards zero by the influence of the prior. This is to be expected and is not surprising. However, it is perhaps less obvious what this means within the simplex: even if 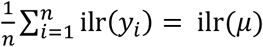, the posterior mode is drawn away from *μ* and towards the centroid of the simplex. More precisely, the posterior mode will be of the form 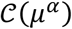 where 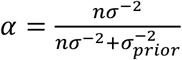 denotes the closure operator which divides a vector by the sum of its components, and the power operation is applied to each component of *μ*. In particular, if *i* denotes the index of the largest component of *μ*, then it can be seen that 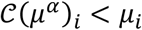. So the largest component of the parameter vector *μ* will be drawn downwards, especially when the number of data points is very small.

The following example illustrates the model in the case *D* = 3, with *n* = 1 representing a scenario where the available data are very limited. Data are simulated from the model 100000 times. For each dataset, an estimate of *μ* is obtained. These estimates are plotted in Figure A1 and Figure A2. There is clearly a tendency for the largest component of of *μ* to be drawn towards the centroid of the simplex.

**Figure A1.**
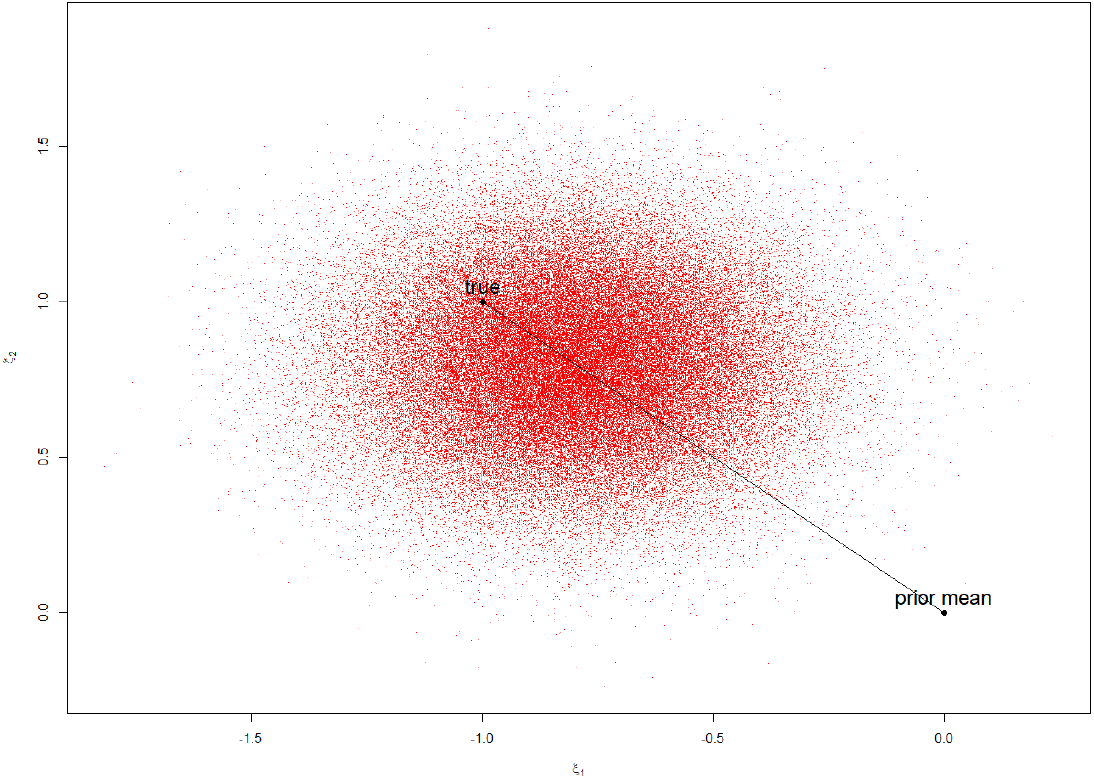
Plot of 100000 estimates (posterior mode) of *μ* in real space. The true value of *μ* is denoted by ‘true’. The straight line connects the true value of *μ* to the prior mean.

**Figure A2.**
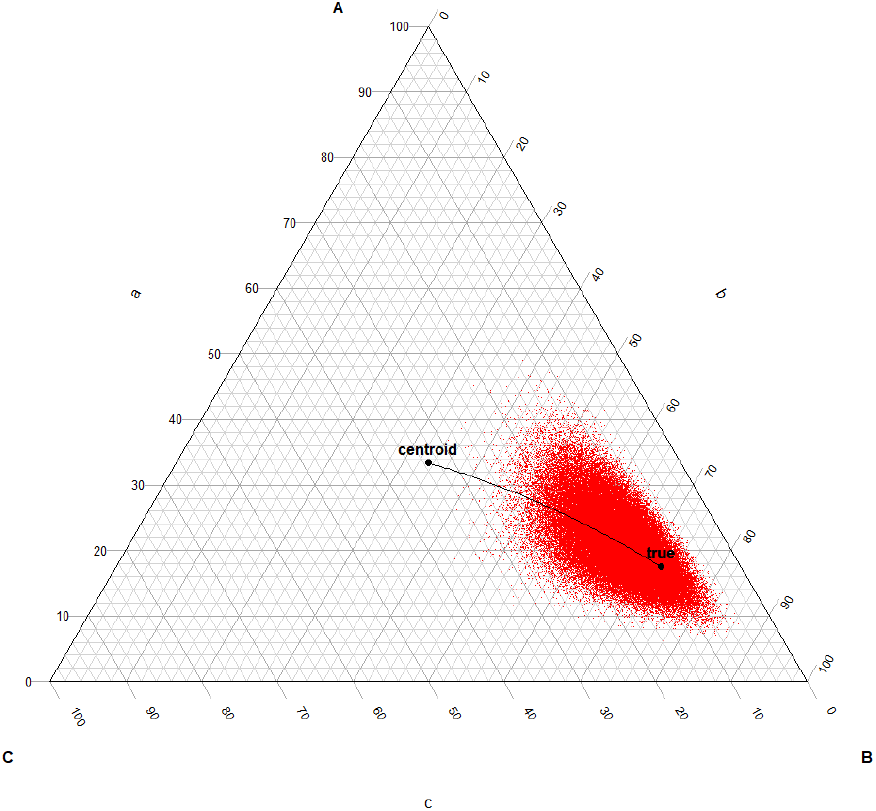
Plot of 100000 estimates (posterior mode) of *μ* in the simplex. The true value of *μ* is denoted by ‘true’. The straight line (in terms of the simplex geometry) connects the true value of *μ* to the centroid of the simplex.

The model setup here is artificial and we have a made a number of unrealistic assumptions. Nevertheless, it is hoped that this shows why it is common that the largest component of a simplex-valued parameter tends to be underestimated.

#### The disease transmission model

We present here an example of inference based on data simulated from the disease transmission model presented in the manuscript, except that the dimension *D* of the simplex parameters has been reduced to *D* = 3. This is achieved by amalgamating some of the land use categories. The reason for doing this is to make it easier to visualise the results of the inference. However, it should be borne in mind that reducing *D* may also reduce some of the previously discussed effects on parameter estimation.

Figures A3 and A4 show the infectivity parameter posterior samples obtained following inference from a set of simulated data with *D* = 3. Comparing the two figures shows that, while of value, the univariate credible intervals do not give a complete picture of the posterior distribution.

**Figure A3.**
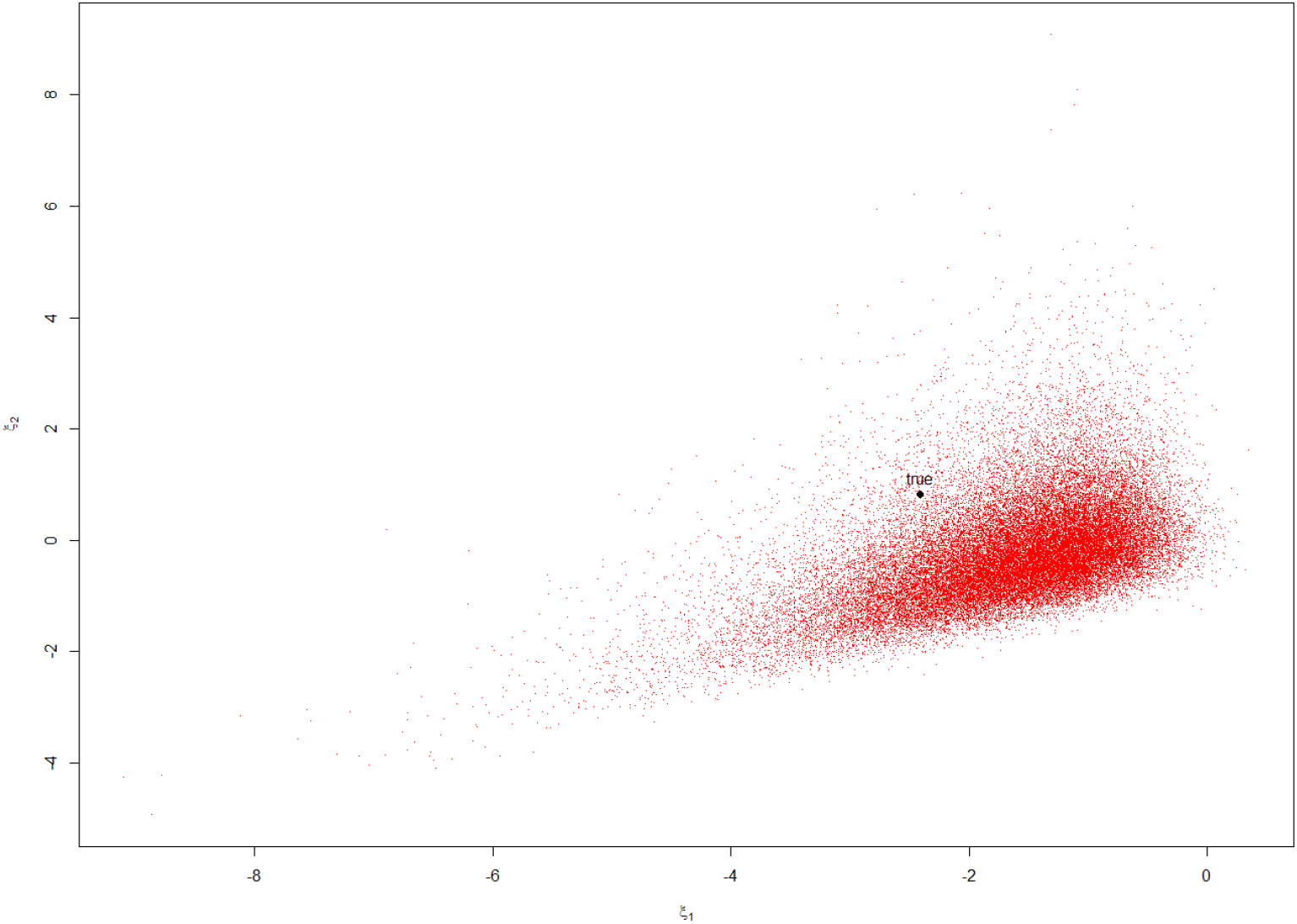
Posterior samples of the infectivity vector in real space, plotted along with the true value. Both univariate 95% credible intervals contain the true parameter.

**Figure A4.**
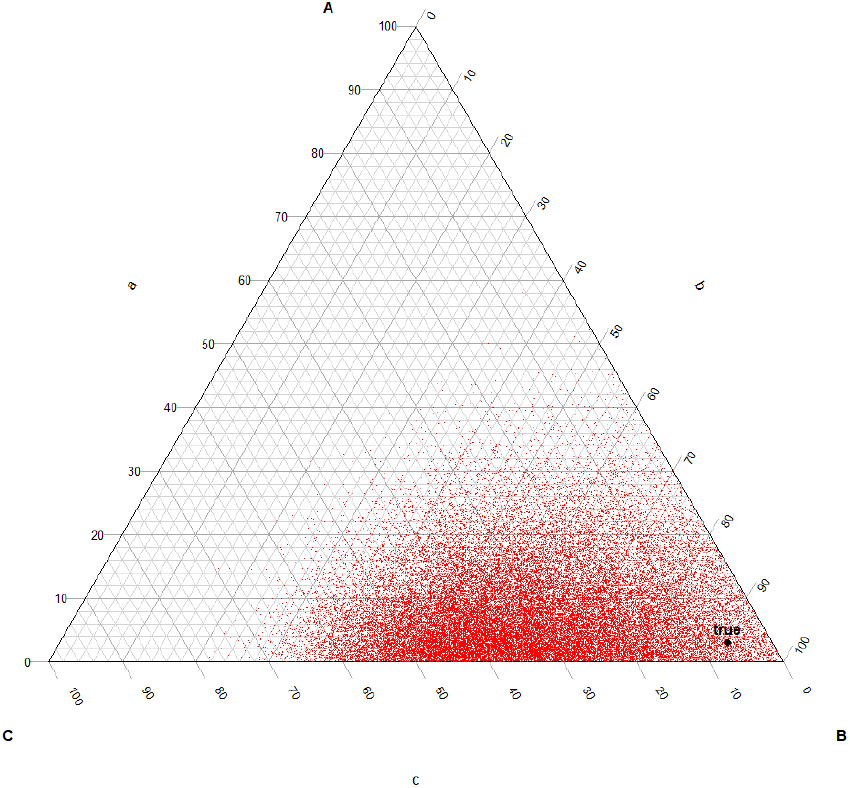
Posterior samples of the infectivity vector in the simplex, plotted along with the true value. The univariate 95% credible intervals for A and C contain the corresponding true values, while the 95% credible interval for B (0.35, 0.89) does not.

Next, we present corresponding results for the disease transmission model with *D* = 6 (Figures A5, A6). In this case, we cannot visualise all dimensions simultaneously. Nevertheless, it appears that the true parameter value is better estimated when viewed in real space than when viewed in simplex space.

**Figure A5.**
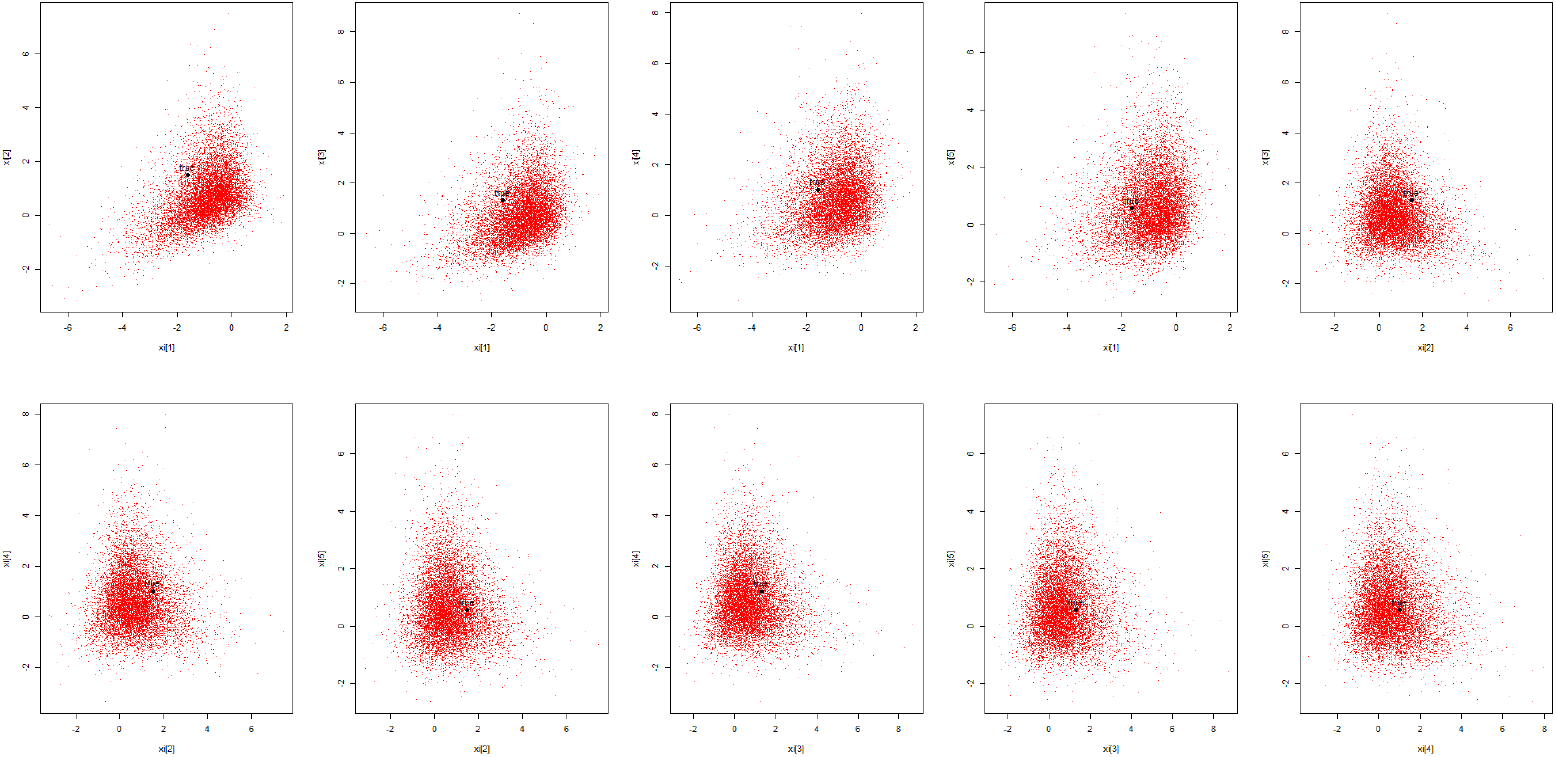
Posterior samples of the infectivity vector in real space, plotted along with the true value. All univariate 95% credible intervals contain the true parameter.

**Figure A6.**
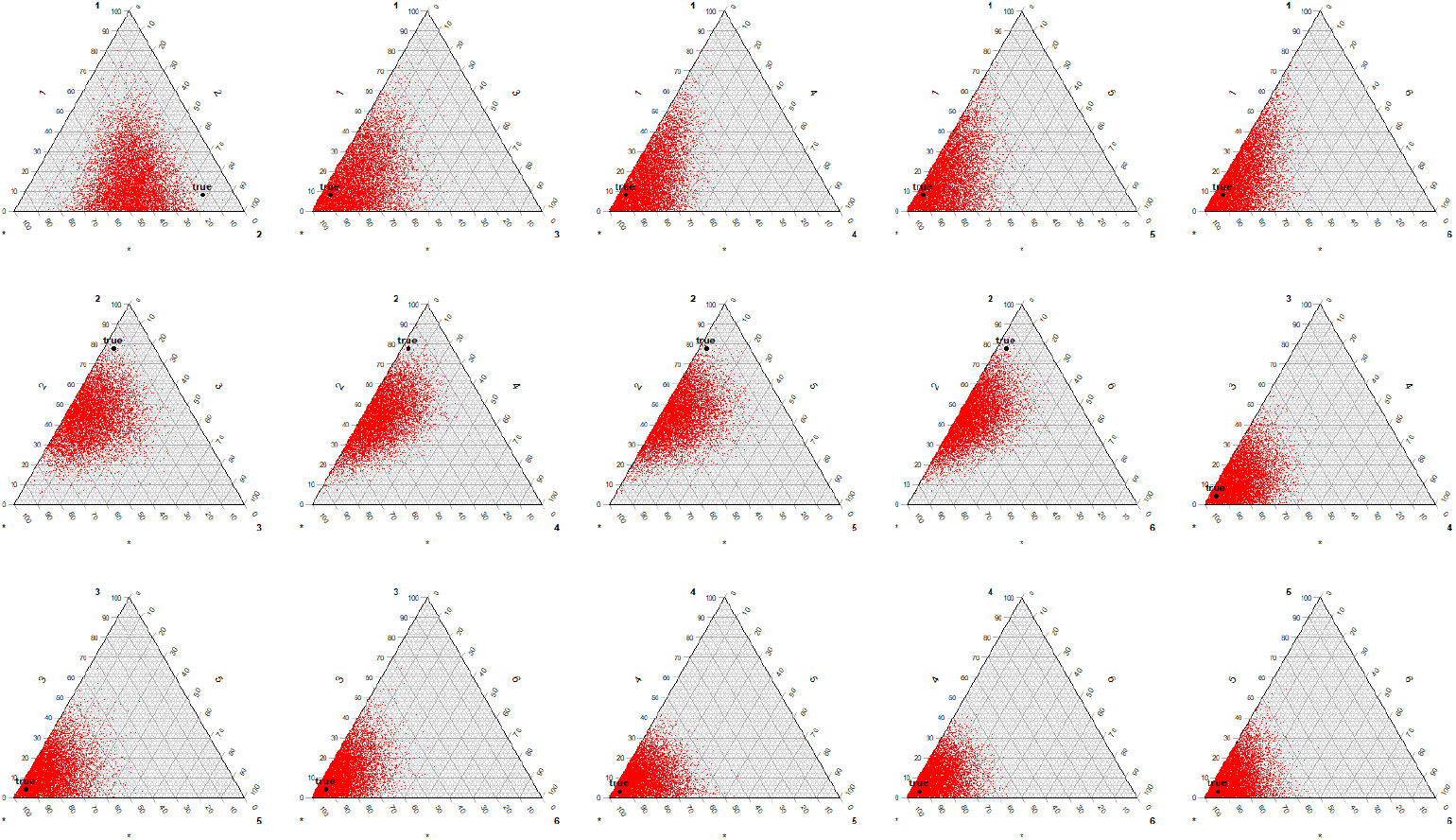
Posterior samples of the infectivity vector in the simplex, plotted along with the true value. The univariate 95% credible intervals all contain the true parameter value, with the exception of parameter 2.

## Appendix 6

### additional figures for Section 3.2

This Appendix provides additional figures for the simulation study in Section 3.2. The plots show true versus inferred parameters for *ξ* = ilr(σ)∈ℝ^5^ and *η* = ilr(γ)∈ℝ^5^. Section 2.2.4 discusses the reason for using the ilr transform on these parameters.

**Figure A7.**
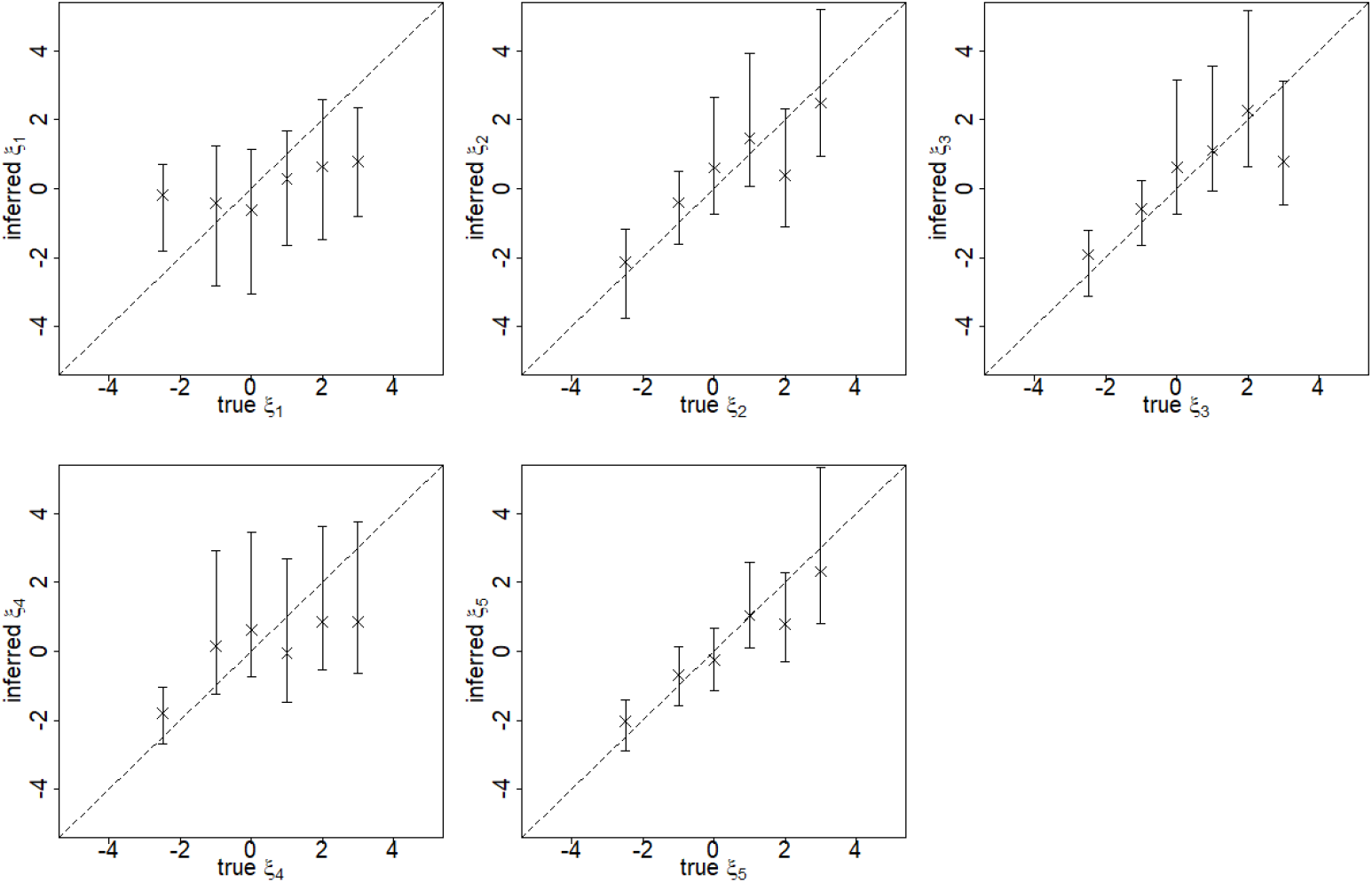
Assessment of the reliability of parameter inference for the constant-in-time iPAR model. Each panel corresponds to a parameter of the model. The ‘true’ value of the parameter is plotted against the posterior median (cross) and posterior 95% credible interval (vertical line). For comparison purposes we also superimpose the diagonal line representing perfect agreement between inferred and true values. This figure includes panels for all five components of *ξ*, the ilr transformed susceptibility parameter.

**Figure A8.**
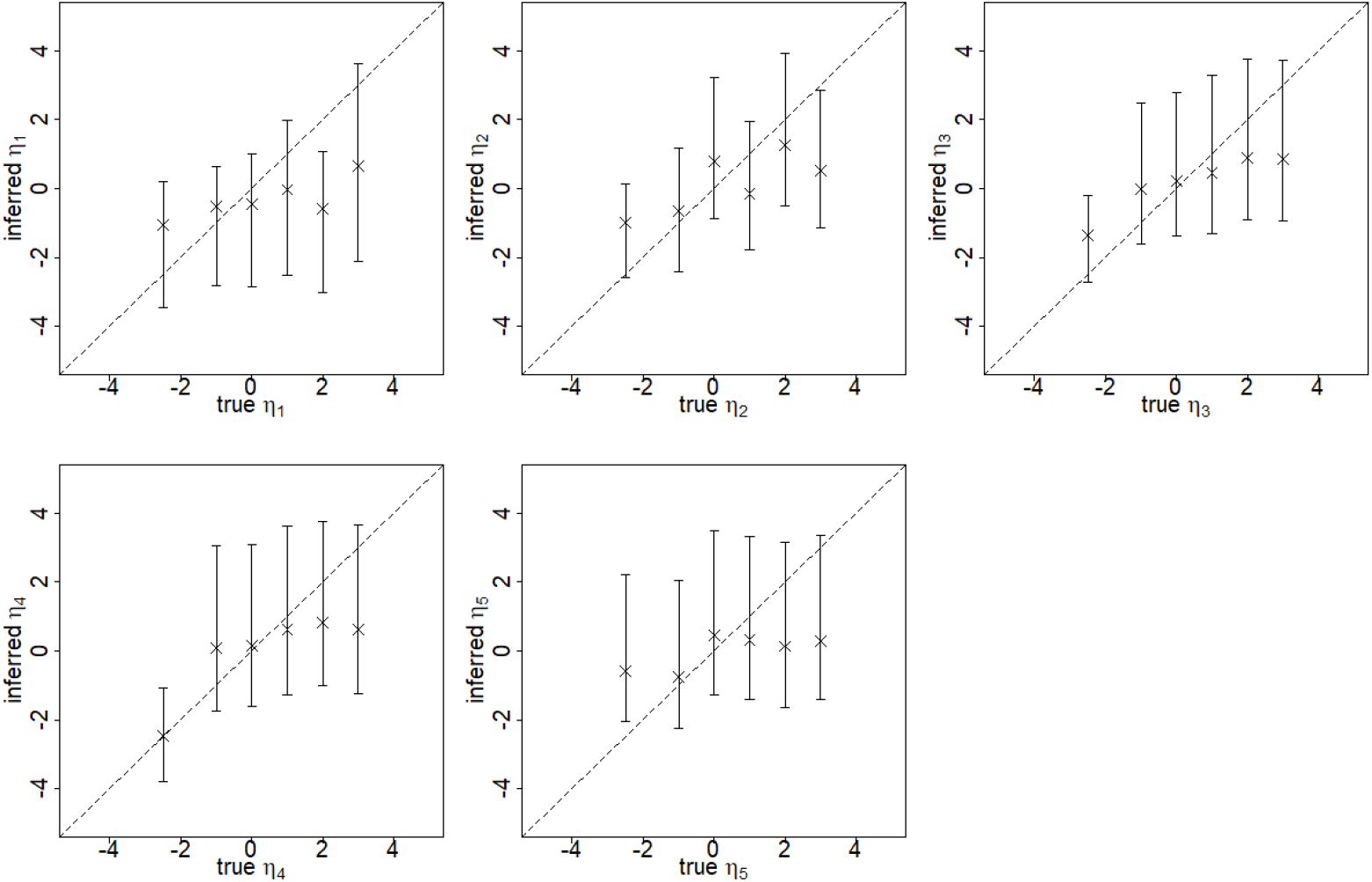
Assessment of the reliability of parameter inference for the constant-in-time iPAR model. Each panel corresponds to a parameter of the model. The ‘true’ value of the parameter is plotted against the posterior median (cross) and posterior 95% credible interval (vertical line). For comparison purposes we also superimpose the diagonal line representing perfect agreement between inferred and true values. This figure includes panels for all five components of *η*, the ilr transformed infectivity parameter.

## Appendix 7

### overestimation of the transmission rate *ρ*

In the simulation studies described in the main text, it was noted that there is typically slight overestimation of the ‘overall’ rate of infection *ρ*, which is proportional to the rate of transmission between two uniform patches (patches containing equal proportions of all land uses). Although we cannot make definitive conclusions regarding why this is the case for any specific model fit, we provide here some intuition as to why there might be overestimation.

First, it might seem surprising that an ‘overall’ rate of transmission could be difficult to estimate. However, it should be borne in mind that *ρ* can be interpreted as the rate of transmission between two uniform patches. If some land uses are not well represented in the covariate data then there will be significant uncertainty in our knowledge of the rate of transmission between two uniform patches. A second issue is that the model aims to disentangle susceptibility and infectivity effects even with relatively poor data (disease cases have imprecise spatial and temporal coordinates). Given that it is known to be particularly difficult to estimate infectivity e.g. see Section 3.4 of the main text, and also Pooley et al. (2020), it is perhaps not surprising that estimation of *ρ* is less straightforward than it might appear.

To be more precise, consider the rate of transmission between an infectious patch *j* and a susceptible patch *i* in the constant-in-time model.

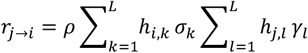

Here, *L* = 6 for the models described in the main text, and we have ignored background transmission (which usually occurs only at a very low level) and the distance dependence (which usually seems well estimated). We further assume that the susceptibility parameter *σ* is well estimated and that the infectivity *γ* is poorly estimated and hence is drawn towards its prior. The outbreak data directly inform us about the rates *r*_*j*→*i*_. So, we might expect overestimation of *ρ* – compensating for the poor estimation of *γ* – when

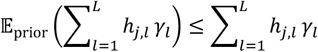

is satisfied on average across all patches *j*. The prior distribution of *γ* is uniform on the simplex and its components sum to 1, so the above condition, when averaged over all *j*, can be rewritten as

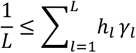

Here, *h*_*l*_ is the proportion of the modelled region (Estonia) occupied by land use category *l*. This condition essentially states that the land uses with highest infectivity are also the most common across the modelled landscape.

In the simulations of Sections 3.3 and 3.4, this condition is always satisfied, and *ρ* is consistently overestimated. In the simulations of Section 3.2, the situation is more complex, with both overestimation and underestimation of *ρ*. We calculated 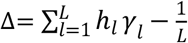 for each of the 78 parameter combinations used in this Section. Low/negative values of Δ seem to be strongly associated with underestimation of *ρ* (Figure A9), which is in line with the informal argument presented above.

**Figure A9.**
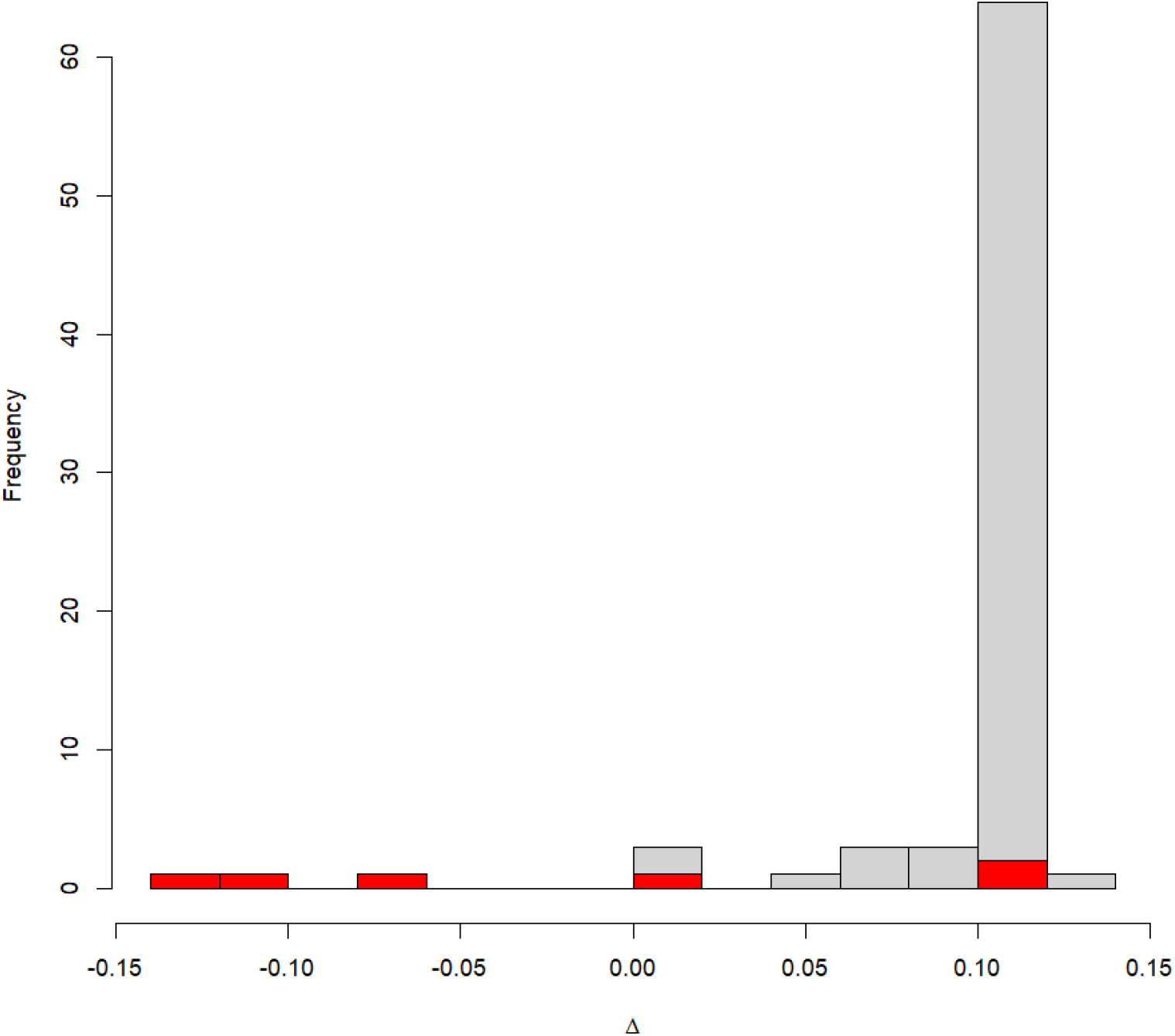
Histogram of Δ values for the 78 simulations (parameter combinations) in Section 3.2. Grey denotes a simulation in which *ρ* is overestimated, red denotes a simulation in which *ρ* is underestimated. Note that the tall peak towards the right simply reflects the way in which the parameter combinations were selected; it does not have any other significance.

## Appendix 8

### details of the setup for simulations in Section 3.4

This appendix details the parameter combinations used in the simulations for Section 3.4 of the main text. Four parameter combinations were used, corresponding to the four coloured rows in Table 1 of the main text. Either a wide or a moderate kernel was employed, together with a choice of a high susceptibility land use and a high infectivity land use. The high susceptibility land use was always ‘broadleaf’, so a parameter combination is defined in terms of its kernel and choice of high infectivity land use. The actual parameter combinations are shown below.

| parameter | Kernel=wide<br>Infe=broadleaf | Kernel=wide<br>Infe=agri | Kernel=moderate<br>Infe=broadleaf | Kernel=moderate<br>Infe=agri |
| --- | --- | --- | --- | --- |
| $\sigma_{\text{urban}}$ | 0.02 | 0.02 | 0.02 | 0.02 |
| $\sigma_{\text{agri}}$ | 0.02 | 0.02 | 0.02 | 0.02 |
| $\sigma_{\text{broadleaf}}$ | 0.90 | 0.90 | 0.90 | 0.90 |
| $\sigma_{\text{conifer}}$ | 0.02 | 0.02 | 0.02 | 0.02 |
| $\sigma_{\text{semi-n}}$ | 0.02 | 0.02 | 0.02 | 0.02 |
| $\sigma_{\text{wetlands}}$ | 0.02 | 0.02 | 0.02 | 0.02 |
| $\gamma_{\text{urban}}$ | 0.02 | 0.02 | 0.02 | 0.02 |
| $\gamma_{\text{agri}}$ | 0.02 | 0.90 | 0.02 | 0.90 |
| $\gamma_{\text{broadleaf}}$ | 0.90 | 0.02 | 0.90 | 0.02 |
| $\gamma_{\text{conifer}}$ | 0.02 | 0.02 | 0.02 | 0.02 |
| $\gamma_{\text{semi-n}}$ | 0.02 | 0.02 | 0.02 | 0.02 |
| $\gamma_{\text{wetlands}}$ | 0.02 | 0.02 | 0.02 | 0.02 |
| $\lambda$ | 0.34 | 0.34 | 0.67 | 0.67 |
| $\rho$ | 7.50 | 7.50 | 7.50 | 7.50 |
| $\epsilon$ | 0.00 | 0.00 | 0.00 | 0.00 |

## Appendix 9

### spread of ASF in Estonia at different spatial scales

In the case study in the main manuscript, we model the spread of ASF in wild boar in Estonia at the 10km resolution. That is, the patches in the model are 10km squares. Here, we briefly explore modelling the same set of outbreak data but at a different, higher spatial resolution of 5km. It is not immediately obvious how the parameter estimates obtained at different resolutions might be related, especially the susceptibility and infectivity estimates for the land use classes in the land use data. The land use covariates, obtained by aggregating over each patch, could be very different at different spatial resolutions. For example, the landscape might seem homogeneous at a coarse resolution – in the sense that each patch has the same land use composition – but at a finer resolution there may be a much higher degree of between-patch variation in land use composition.

#### 5km spatial resolution: spatially aggregated outbreak data

First, we explore the spatially aggregated outbreak data at different points in time. Figures A10, A11, A12 show how the infection spreads across Estonia, at the 5km resolution. These figures are analogous to Figure 6 in the main manuscript, but at this higher spatial resolution. There is clear visual evidence of distance-dependence in the spread of disease.

**Figure A10.**
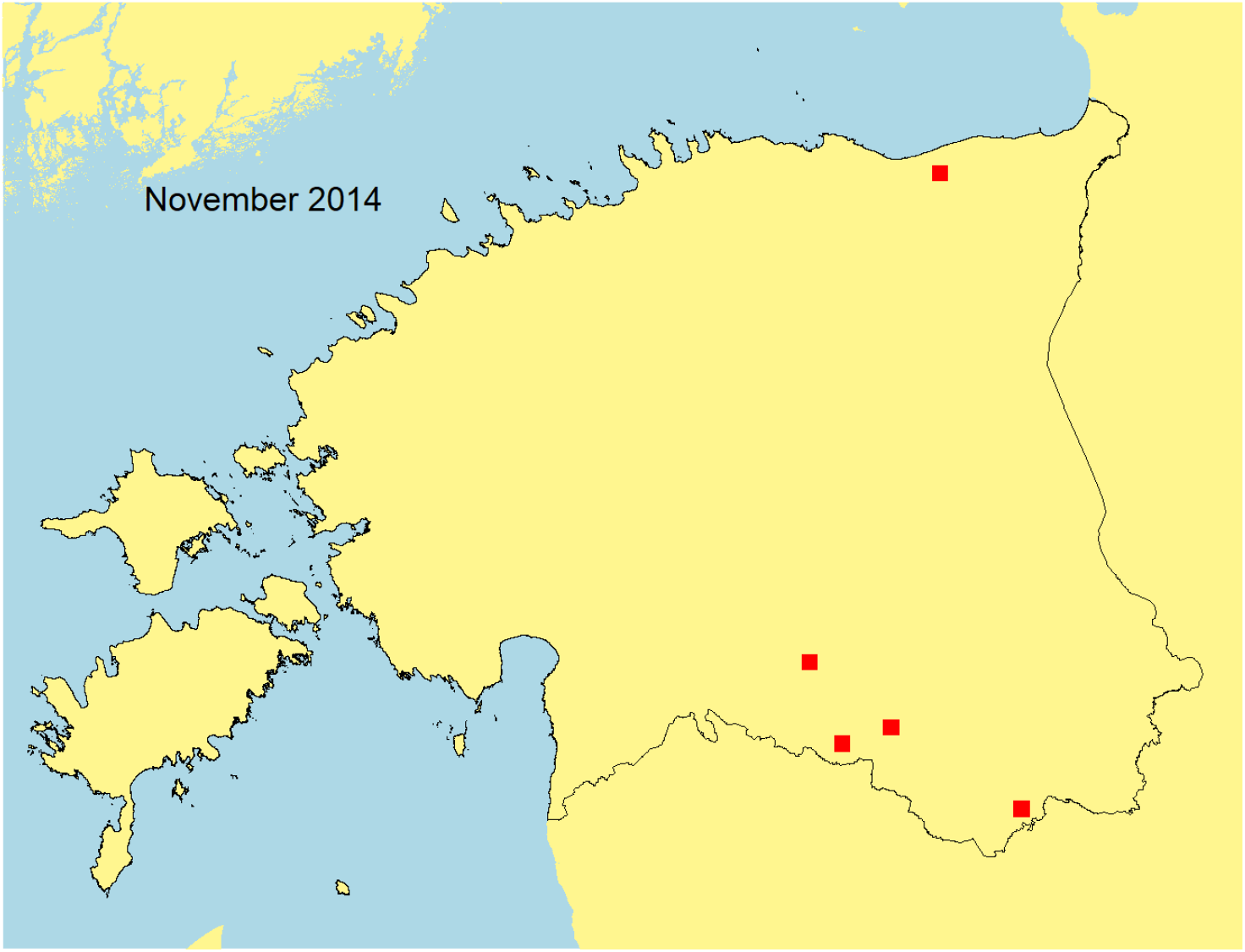
Case reports: Spatially aggregated (5km resolution) African Swine Fever wild boar cases reported in Estonia. Snapshot as at November 2014.

**Figure A11.**
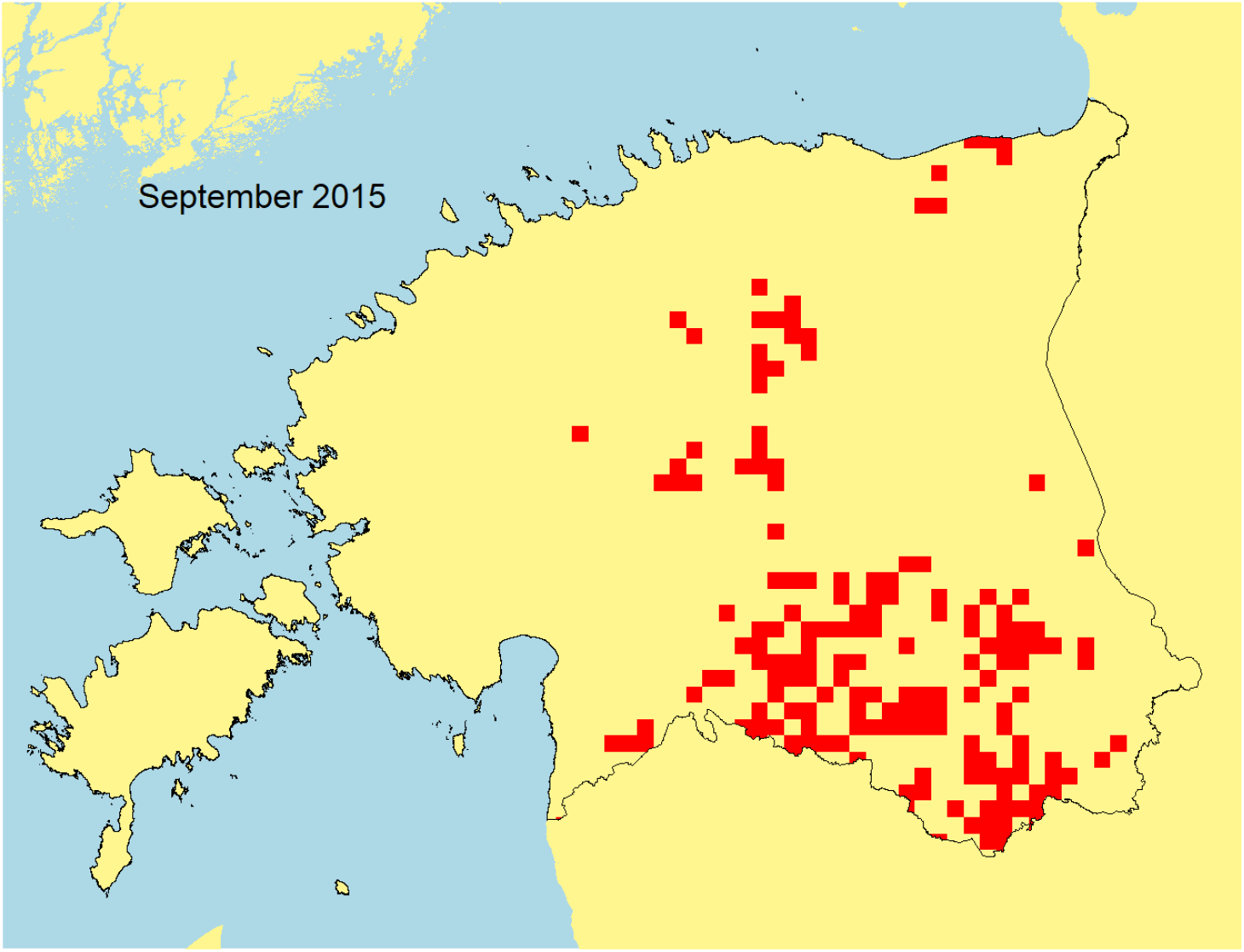
Case reports: Spatially aggregated (5km resolution) African Swine Fever wild boar cases reported in Estonia. Snapshot as at September 2015.

**Figure A12.**
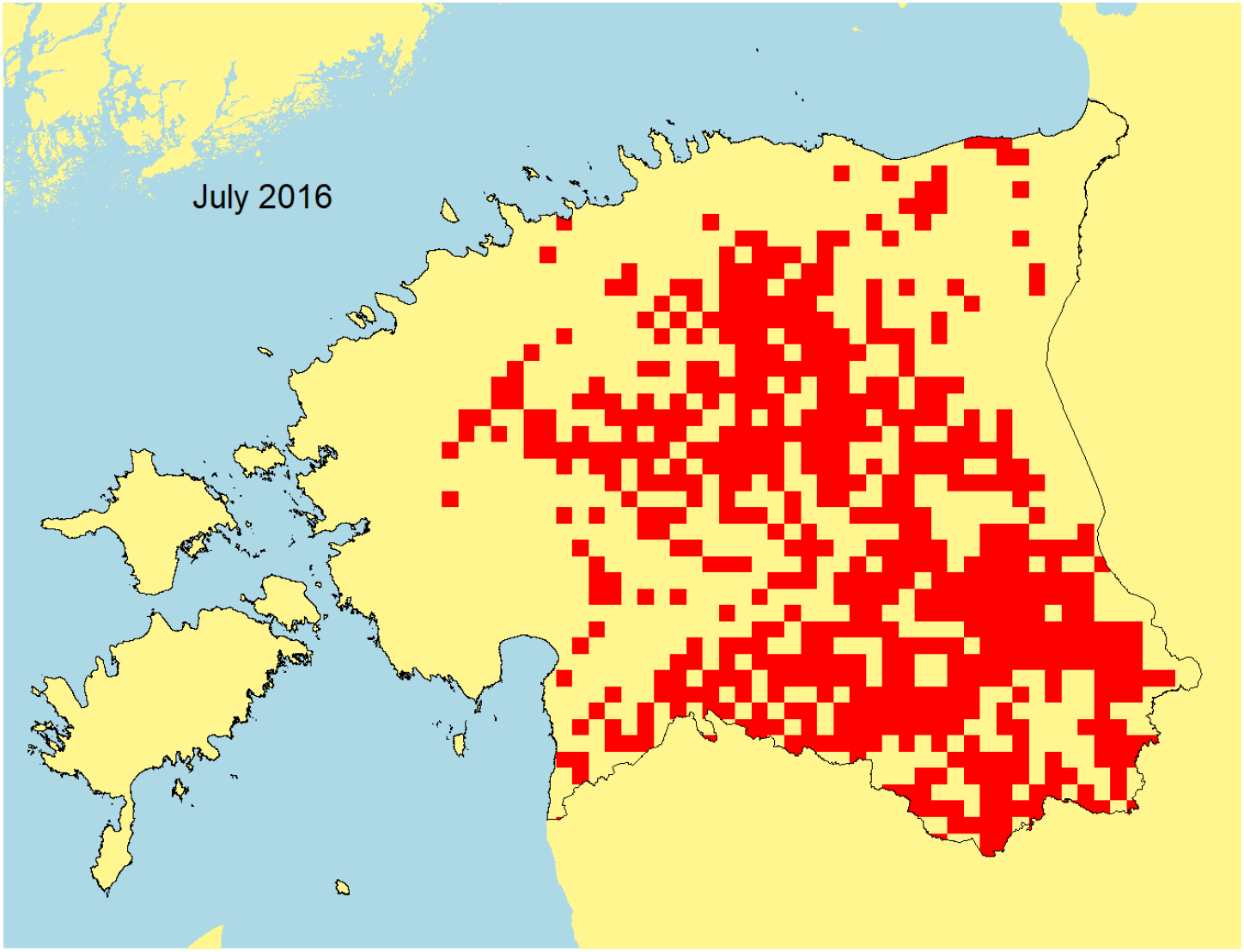
Case reports: Spatially aggregated (5km resolution) African Swine Fever wild boar cases reported in Estonia. Snapshot as at July 2016.

#### 5km spatial resolution: model parameter estimates and interpretation

We fitted the constant-in-time model to the same set of outbreak data as used in the case study in the main text, but at the 5km spatial resolution. The structure of the model is the same as that used in the case study in the main text, except for the higher spatial resolution. Parameter estimates from the fitted model are shown below.

|  | Estonia constant in time |  |  |
| --- | --- | --- | --- |
| parameter | lower | median | upper |
| $\sigma_{\text{urban}}$ | 0.00 | **0.01** | 0.06 |
| $\sigma_{\text{agri}}$ | 0.19 | **0.26** | 0.34 |
| $\sigma_{\text{broadleaf}}$ | 0.20 | **0.28** | 0.37 |
| $\sigma_{\text{conifer}}$ | 0.13 | **0.21** | 0.29 |
| $\sigma_{\text{semi-n}}$ | 0.05 | **0.17** | 0.29 |
| $\sigma_{\text{wetlands}}$ | 0.01 | **0.05** | 0.10 |
| $\gamma_{\text{urban}}$ | 0.00 | **0.08** | 0.34 |
| $\gamma_{\text{agri}}$ | 0.28 | **0.50** | 0.74 |
| $\gamma_{\text{broadleaf}}$ | 0.01 | **0.13** | 0.34 |
| $\gamma_{\text{conifer}}$ | 0.00 | **0.06** | 0.20 |
| $\gamma_{\text{semi-n}}$ | 0.00 | **0.05** | 0.23 |
| $\gamma_{\text{wetlands}}$ | 0.00 | **0.09** | 0.31 |
| $\lambda$ | 1.60 | **1.87** | 2.30 |
| $\rho$ | 0.57 | **0.76** | 1.09 |
| $1000\varepsilon$ | 27.56 | **33.32** | 39.64 |

When interpreting the above parameter estimates, it is worth bearing in mind that the estimates may not necessarily match those obtained at the 10km resolution. The pattern of disease spread may appear quite different at different spatial resolutions. Nevertheless, the susceptibility and infectivity estimates are broadly similar to those at the 10km resolution, as can be seen by comparing the table above with Table 4 in the main text. Broadleaf woodland remains the land use with the highest susceptibility, while agricultural land remains the land use with the highest infectivity.

Regarding estimates of the transmission kernel parameter *λ*, it is worth remembering that distance is defined in terms of the patch dimensions, so when the patch size is reduced we also change the scale on which distance is measured. The power law transmission kernel is scale invariant, so we might perhaps expect the estimates of *λ* to be similar at different spatial scales. On the other hand, the transmission is unlikely to exactly follow a power law kernel. What we actually find is that *λ* is estimated to be a little higher at 5km resolution, suggesting something a little closer to ‘nearest neighbour’ transmission between patches. The overall rate *ρ* is also lower, while the background transmission rate *ε* is higher. In summary, background transmission seems relatively more important when the data are viewed at the 5km resolution.

## Appendix 10

### Persistence of infection analysis

In this section, we take a closer look at the case report data for ASF in wild boar in Estonia. The assumed disease transmission model is a form of spatial SI model, so once a patch becomes infected it remains infected. By looking at the local persistence of ASF infection within each modelled 10km square patch, we can assess how realistic the SI model infection persistence assumption actually is. Within each patch there may be one of more disease case reports. We computed the difference between the first and last disease report time for each patch covering Estonia. A histogram of these differences is shown in Figure A13. We only include patches with reported cases before June 2015 because it is only these patches, infected during the early stages of the outbreak, for which long term persistence of infection can be fully assessed. Infection is usually persistent for at least a year, and often up to four years. The observed persistence of ASF infection in wild boar goes some way to support the SI infection persistence assumption used in the transmission model.

**Figure A13.**
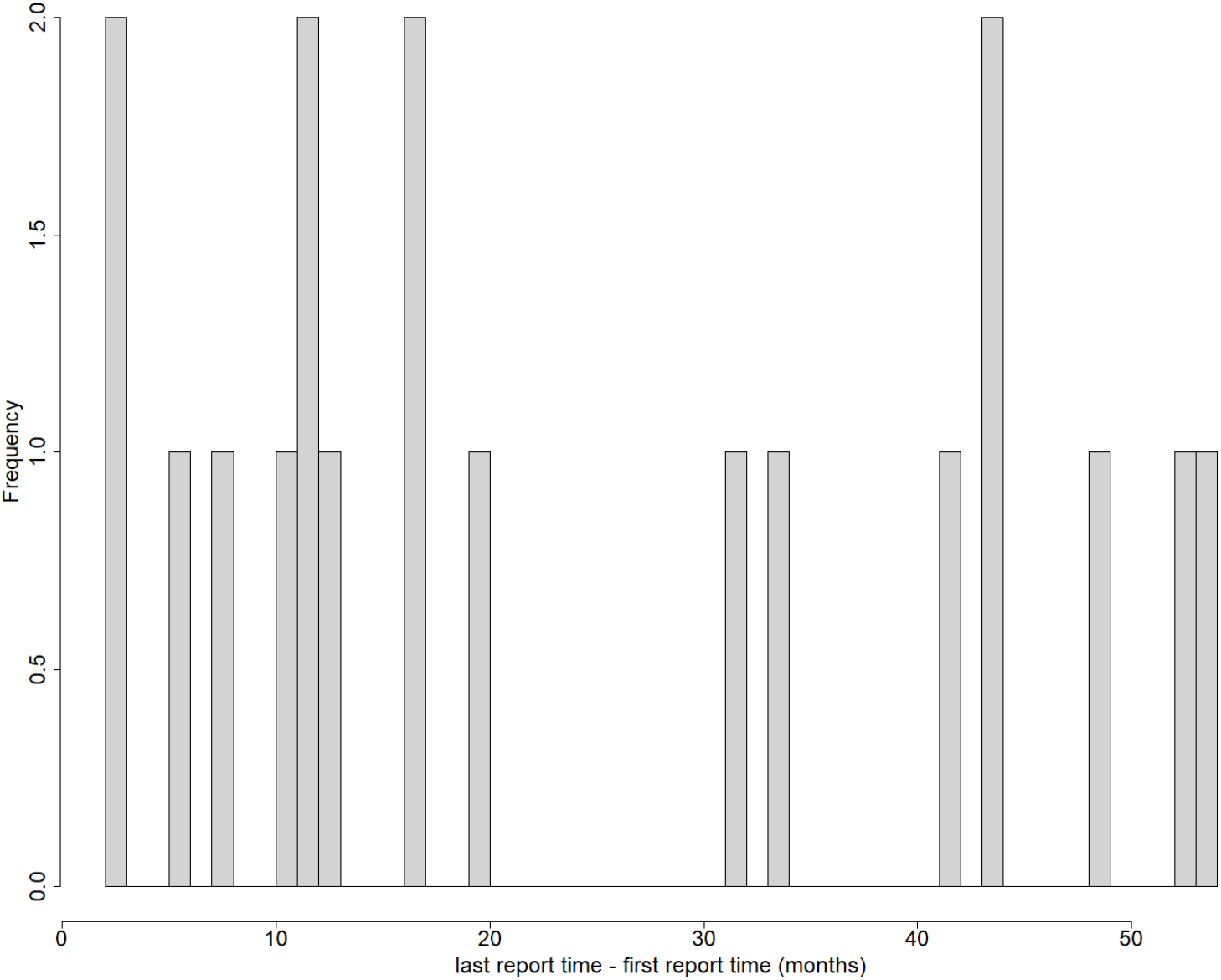
Persistence of infection in 10km square patches covering Estonia. A histogram of the difference between the first disease report time and the last disease report time, as computed for each patch covering Estonia. Patches are excluded if there were no reported cases before June 2015.

One limitation of this analysis is that a relatively small number of patches were infected early on in the outbreak. To gain a broader picture, we also investigated outbreak data from neighbouring countries in the Baltic states – Latvia and Lithuania. Figure A14 shows the same analysis, but conducted over all three countries in the Baltic states. Here, there is even stronger evidence for the long-term persistence of infection in wild boar. Infection usually persists for over 2 years, frequently for 4 or 5 years. This analysis further supports the SI infection persistence assumption used in the transmission model.

**Figure A14.**
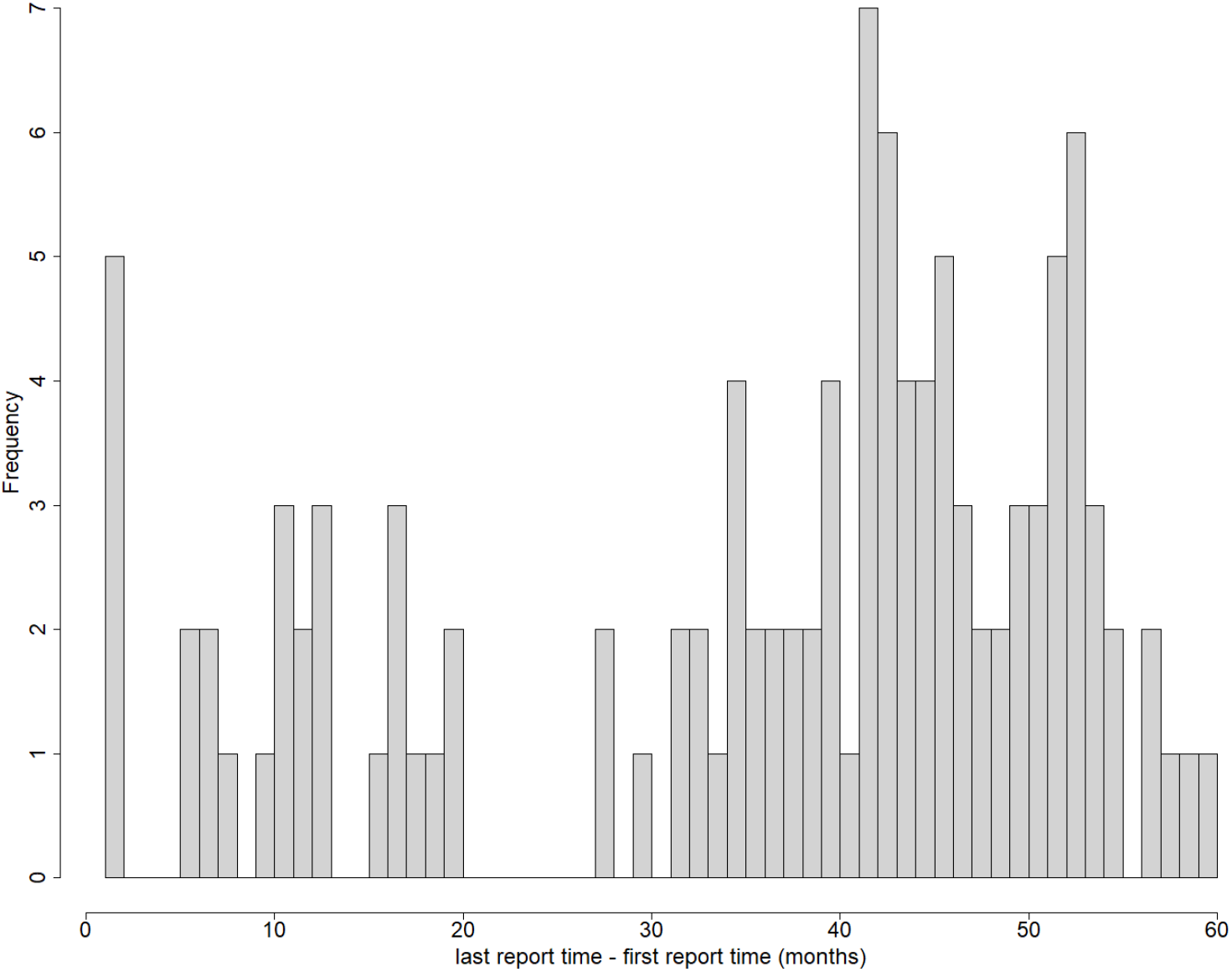
Persistence of infection in 10km square patches covering the Baltic states – Estonia, Latvia and Lithuania. A histogram of the difference between the first disease report time and the last disease report time, as computed for each patch covering the Baltic states. Patches are excluded if there were no reported cases before June 2015.

## Appendix 11

### Narrow transmission kernel simulations

This section is motivated by the predictive performance results from the case study, as shown in Table 5 in the main text. The homogeneous and heterogeneous models give very similar predictive performances. Moreover, the predictions seem much better overall than those obtained in Section 3.4 – compare Table 5 with Table 1. To help explain these results, we show here the results of simulations similar to those in Section 3.4 but with (a) non-kernel parameters close to those estimated from the real data (Table 4), and (b) the kernel parameter set to give various narrow kernels. The precise parameter combinations used in these simulations are shown in the table below. Three distinct parameter combinations were used.

| parameter | Narrow kernel | Very narrow kernel | Extremely narrow kernel |
| --- | --- | --- | --- |
| $\sigma_{\text{urban}}$ | 0.01 | 0.01 | 0.01 |
| $\sigma_{\text{agri}}$ | 0.09 | 0.09 | 0.09 |
| $\sigma_{\text{broadleaf}}$ | 0.42 | 0.42 | 0.42 |
| $\sigma_{\text{conifer}}$ | 0.20 | 0.20 | 0.20 |
| $\sigma_{\text{semi-n}}$ | 0.20 | 0.20 | 0.20 |
| $\sigma_{\text{wetlands}}$ | 0.07 | 0.07 | 0.07 |
| $\gamma_{\text{urban}}$ | 0.08 | 0.08 | 0.08 |
| $\gamma_{\text{agri}}$ | 0.82 | 0.82 | 0.82 |
| $\gamma_{\text{broadleaf}}$ | 0.03 | 0.03 | 0.03 |
| $\gamma_{\text{conifer}}$ | 0.02 | 0.02 | 0.02 |
| $\gamma_{\text{semi-n}}$ | 0.03 | 0.03 | 0.03 |
| $\gamma_{\text{wetlands}}$ | 0.03 | 0.03 | 0.03 |
| $\lambda$ | 1.45 | 3.00 | 5.00 |
| $\rho$ | 4.54 | 4.54 | 4.54 |
| 1000ε | 7.65 | 7.65 | 7.65 |

The predictive performance obtained in these simulations is shown in the table below. The results obtained are similar to those obtained in Table 5 in the case study i.e. very high discriminatory power, and close agreement between the heterogeneous and homogeneous fitted models.

| | | Discrimination metrics | | $N_{est}/N_{truth}$ | |
| --- | --- | --- | --- | --- | --- |
| Kernel | Model | mean AUC | mean TPR @ 5% FPR | reliability (%) | mean bias |
| Narrow | homog | 0.77 | 0.22 | 95 | 0.02 |
|  | heter | 0.77 | 0.23 | 100 | 0.01 |
| Very narrow | homog | 0.84 | 0.37 | 95 | 0.04 |
|  | heter | 0.85 | 0.39 | 100 | 0.02 |
| Extremely narrow | homog | 0.86 | 0.36 | 95 | 0.04 |
|  | heter | 0.86 | 0.38 | 100 | 0.02 |

